# mRNA 3′UTRs chaperone intrinsically disordered regions to control protein activity

**DOI:** 10.1101/2025.07.02.662873

**Authors:** Yang Luo, Yaofeng Zhong, Sudipto Basu, Ming-Chung Wu, Christine Mayr

**Affiliations:** Cancer Biology and Genetics Program, Sloan Kettering Institute, New York, NY 10065, USA; Biochemistry, Cell, and Molecular Biology (BCMB) Allied Program, Weill Cornell Graduate School of Medical Sciences, New York, NY 10065, USA

**Keywords:** 3′UTR, intrinsically disordered regions, protein folding, cotranslational folding, transcription factor, chromatin regulator, RNA-IDR interaction, RNA-based molecular chaperone, RNA-mediated protein folding, biomolecular condensates

## Abstract

Nearly 3,000 human mRNA 3′UTRs have hundreds of highly conserved nucleotides, but their biological roles are unclear. These mRNAs mostly encode proteins with long intrinsically disordered regions (IDRs), including MYC, UTX, and JMJD3. We show that these proteins are only fully active when translated from mRNA templates that include their 3′UTRs, raising the possibility of functional interactions between 3′UTRs and IDRs. Rather than affecting protein abundance or localization, we find that the *KDM6B* 3′UTR in the mRNA template changes the folding of the encoded IDR-containing JMJD3 protein. It promotes IDR-IDR interactions and suppresses folding between domains, suggesting that RNA acts as IDR chaperone that prevents interference of hydrophobic clusters in the IDR with folding of the structured domain. mRNA-based IDR chaperones are enriched in meshlike cytoplasmic condensates, suggesting localized chaperone activity. As hydrophobic clusters in IDRs are widespread, our data suggest that 3′UTR-dependent protein folding could be a widely used mechanism for activity regulation of transcriptional regulators.

## Introduction

Messenger RNAs (mRNAs) are the templates for protein synthesis which makes them central components of gene expression regulation. Each mRNA contains a coding sequence (CDS), 5′ untranslated region (UTR), and 3′UTR. Transcriptome-wide mRNA 3′ end sequencing in human primary cells revealed that overall, CDS and 3′UTRs both contribute nearly 50% of mRNA sequence^1,2^. Whereas the CDS is translated into the protein sequence, the biological roles of 3′UTRs are less clear.

Already 20 years ago, it was noticed that a subset of human 3′UTRs have regions of hundreds of nucleotides of exceptionally highly sequence conservation, but their functional relevance is unknown^3^. More recently, a relationship between 3′UTR sequence conservation, nucleotide content, and mRNA destabilization was recognized, where rapidly evolving 3′UTRs are GC-rich and have destabilizing effects on mRNAs, whereas conserved 3′UTRs are GC-poor and lack destabilizing function^4^.

Individual 3′UTRs have been shown to regulate mRNA stability, mRNA localization, and protein localization^5–8^. However, they often act in a context-specific manner, where the presence of a regulatory element does not guarantee a functional effect^9,10^. Moreover, 3′UTRs play important roles in cytoplasmic organization^11–14^. The view on cytoplasmic organization has substantially evolved within the past few years by the discovery that the cytoplasm is compartmentalized not only by membrane-enclosed organelles, but also by membraneless compartments. Importantly, several research groups found that mRNAs do not distribute randomly within the cytoplasm but rather segregate into membrane-associated, condensate-enriched, and unbound cytosolic mRNAs^13,15–20^. In a few cases, the 3′UTR-determined location of protein translation has been shown to control the functions of the newly synthesized proteins^8,11,21^. However, it is currently unclear, whether translation of an mRNA in cytoplasmic condensates versus the cytosol has broader biological consequences.

During folding of natively structured proteins, hydrophobic residues cluster together to form a hydrophobic core. However, little is known about the folding of large, multidomain proteins^22,23^. Folding of multidomain proteins begins co-translationally with the help of molecular chaperones, while the ribosome still elongates the nascent polypeptide chain^23,24^. In addition to folded domains, most human proteins have intrinsically disordered regions (IDRs), which are protein segments with a lower fraction of hydrophobic amino acids than folded domains^25^. The sequence-structure-function relationships of IDRs remain poorly characterized, as IDRs do not adopt a single stable conformation in isolation but sample a wide array of distinct conformational states^26,27^. Nevertheless, two-thirds of IDRs adopt defined secondary structures when bound to their partners, which can be other proteins or RNA^26,28–32^. The latter pathway could be widespread, supported by the recent discovery of prevalent RNA-IDR interactions in cells^33,34^. As IDRs are strongly enriched in regulatory proteins, including transcription factors and chromatin regulators, a better understanding by which IDRs influence protein function is of broad biological significance^35^.

Here, we show that inclusion of the highly conserved 3′UTRs in mRNA templates encoding MYC, UTX, and JMJD3 is required for full cellular transcriptional or histone demethylase activity of the encoded proteins in the nucleus. For JMJD3, protein activity is not controlled by protein abundance or localization, but by a 3′UTR-dependent change in protein folding, shown by crosslinking mass spectrometry of cellular JMJD3. Altered protein folding is recapitulated in vitro, where RNA-IDR interaction is necessary and sufficient for a change in protein domain interaction preference. RNA-IDR binding promotes IDR-IDR interaction and prevents IDR-folded domain interaction. Our data suggest a model, in which the 3′UTR co-translationally promotes IDR-IDR interaction and suppresses hydrophobic clusters in the IDR from interfering with folding of the JMJD3 structured domain. Based on low-resolution in-cell crosslinking data obtained from ten candidates, we propose that only the subset of 4,910 human proteins whose IDRs contain hydrophobic amino acid clusters are potentially susceptible for 3′UTR-dependent changes in protein folding. For these candidates, we observed that the protein sequence alone is not sufficient to generate a biologically fully active protein but requires regulatory elements in the 3′UTR of their mRNA template.

## Results

### mRNAs with highly conserved 3′UTRs encode proteins with long IDRs

We previously performed transcriptome-wide mapping of 3′UTRs and observed that 3′UTRs make up nearly 50% of human mRNA sequence (Fig. S1A)^1,2^. We noticed that a subset of human 3′UTRs have sequences that are markedly conserved across vertebrate species (Fig. 1A). To identify human 3′UTRs with a large number of conserved nucleotides, we used Phylogenetic P values (PhyloP conservation scores) obtained from 100 vertebrate genomes, which include fish, birds, and mammals^3,36^. We define a 3′UTR as highly conserved (HC), when it has at least 150 nucleotides with PhyloP scores ≥2, only considering regions with at least four consecutive nucleotides passing the threshold. Across the entire human genome only 0.7% of bases meet these criteria. With respect to human 3′UTRs, we detect 2,714 HC 3′UTRs, 7,947 non-conserved (NC) 3′UTRs that have zero conserved nucleotides, given our criteria, and an intermediate group (Fig. 1B, Table S1).

**Figure 1.**
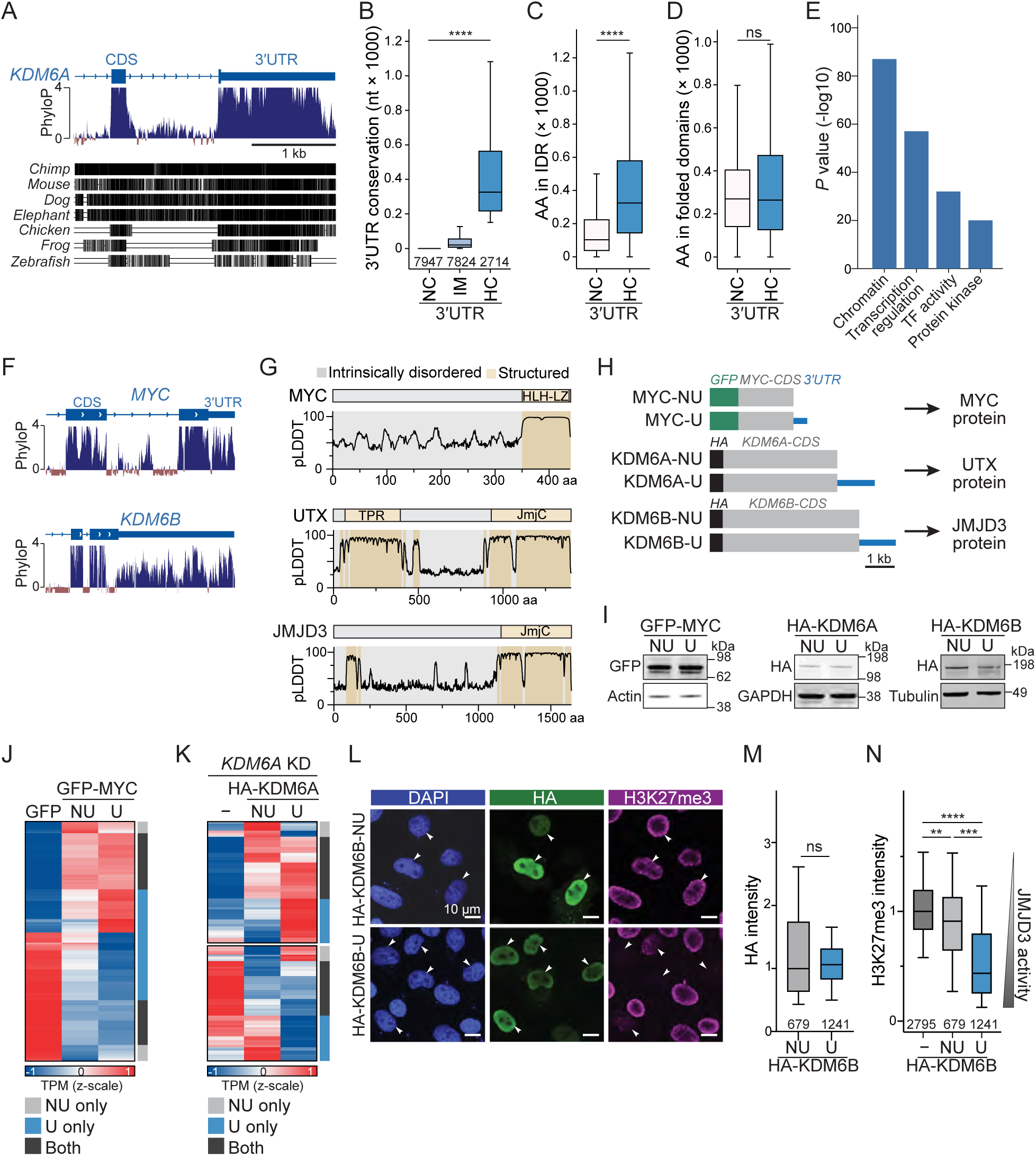
HC 3′UTRs regulate activity of transcription and chromatin regulators independently of protein abundance. **A.** Sequence conservation of 100 vertebrate genomes, obtained from the UCSC PhyloP100way track is shown for the genomic context depicting the last two exons of the *KDM6A* gene. **B.** Classification of human genes based on the number of conserved nucleotides (nt) in human 3′UTRs obtained from PhyloP. Non-conserved (NC), 0 conserved nt (*N* = 7947), intermediate (IM), intermediate, 4-150 nt (*N* = 7824), highly conserved (HC) >150 nt (*N* = 2714). Kruskal-Wallis test, *P* = 0. **C.** Number of residues in the IDRs of proteins encoded by mRNAs shown in (B). IDR if IUPreD2A score ≥ 0.33. AA, amino acid. Mann-Whitney test: *P* = 0. **D.** As in (C), but the number of residues in the folded domains is shown. Folded domain, if IUPreD2A score < 0.33. Mann-Whitney test: ns, not significant. **E.** Gene ontology analysis of mRNAs with HC 3′UTRs. The top four functional gene classes and Benjamini Hochberg-adjusted *P* values are shown. TF, transcription factor. **F.** As in (A) but shown for *MYC* and *KDM6B*. **G.** Prediction of structured domains (pLDDT ≥ 80, brown) and IDRs (pLDDT < 80, gray) using AlphaFold2 pLDDT scores. Names of annotated protein domains are given. **H.** cDNA vector constructs used for protein activity assays in cells, showing the tag, CDS (gray) and full-length 3′UTR (blue). NU, no 3′UTR; U, 3′UTR was included in the vector. **I.** Western blot analysis after transfection of constructs shown in (H) into U2OS, MCF7, and HeLa cells. Actin, GAPDH, and Tubulin were used as loading controls. **J.** Genes induced (red) or repressed (blue) upon transfection of GFP-MYC-U or GFP-MYC-NU compared to GFP control in U2OS cells. Shown are genes with |log2FC| > 0.3 and adjusted *P* value < 0.1, which were used to identify indicated groups by hierarchical clustering. Genes with significant change only in NU vs GFP (NU only; *N* = 185), significant change only in U vs GFP (U only, *N* = 753), and significant change in NU vs GFP and U vs GFP (both, *N* = 686). TPM, transcripts per million. **K.** As in (J) but using MCF7 cells with shRNA-mediated knockdown of endogenous *KDM6A,* which were subsequently transfected with HA-KDM6A-NU or HA-KDM6A-U. Shown are transcripts significantly up- or downregulated by *KDM6A* knockdown relative to control (DESeq2 adjusted *P* value < 0.05). These genes were used for subsequent clustering analysis. Clustering was performed separately for up- and downregulated groups. Shown are all genes, whose expression was at least partially rescued by expression of HA-KDM6A-NU or HA-KDM6A-U. NU only, *N* = 77; U only, *N* = 173; both, *N* = 151. **L.** Immunofluorescence assay to assess 3′UTR-dependent cellular JMJD3 activity in HeLa cells after transfection of HA-KDM6B-NU or HA-KDM6B-U. HeLa cells have low levels of endogenous JMJD3. Nuclei were stained by DAPI. Anti-HA staining (green) shows transfected JMJD3 protein, and a reduction in anti-H3K27me3 staining (magenta) is used to infer JMJD3 cellular activity. White arrowheads mark transfected cells. Representative images are shown. **M.** Quantification of transfected JMJD3 abundance as HA intensity values from (L). Shown are boxplots depicting median, 25^th^ and 75^th^ percentiles and 5-95% confidence intervals from *N* = 3 independent experiments. Mann-Whitney test: ns, not significant. **N.** 3′UTR-dependent JMJD3 cellular activity was measured as a reduction in nuclear H3K27me3 intensity in transfected cells compared to non-transfected cells from the experiment shown in (L). The nuclear H3K27me3 intensity in each cell was normalized to the median intensity of non-transfected cells (HA < 500 units). Shown as in (M) from *N* = 3 independent experiments. Mann-Whitney test, **, *P =* 2.3 x 10^−16^; ***, *P* = 6.3 x 10^−71^; ****, *P =* 5.8 x 10^−252^; ns, not significant.

We noticed that mRNAs with NC 3′UTRs mostly encode small, well-folded proteins, whereas mRNAs with HC 3′UTRs encode proteins with long IDRs (Fig. 1C, 1D). We define IDRs as protein regions with IUPred2A values ≥0.33^37^. We obtained similar results when using AlphaFold-predicted local distance difference test (pLDDT) scores for IDR identification (Fig. S1B)^38^. Gene ontology analysis showed that proteins encoded by mRNAs with HC 3′UTRs are substantially enriched in chromatin regulators and transcription factors (Fig. 1E)^39^, consistent with previous observations that these factors are overrepresented among proteins with long IDRs^35,40^.

### Including 3′UTRs in mRNA templates increases protein activity of transcriptional regulators

To investigate the biological roles of HC 3′UTRs, we selected three candidate genes (*MYC*, *KDM6A, KDM6B*), whose mRNAs have HC 3′UTRs and encode proteins with long IDRs (Fig. 1A, 1F, 1G). We transiently expressed these transcriptional regulators using cDNA expression vectors that either contained the CDS only (called NU, for ‘no 3′UTR’) or that contained the CDS and the corresponding full-length 3′UTR (called U; Fig. 1H). We observed that protein abundance and subcellular localization of the transcriptional regulators were unaffected when they were expressed in the presence or absence of their 3′UTRs (Fig. 1I, S1C, S1D).

Next, we investigated potential differences in protein activity, when the proteins were expressed from NU or U constructs. We expressed MYC-U or MYC-NU in U2OS cells, where endogenous MYC is low, and performed RNA-seq. Compared to the GFP-only control, the vector containing MYC-U induced or repressed nearly twice as many genes as the MYC-NU vector (Fig. 1J, Table S2). This indicates that inclusion of the 3′UTR in the mRNA template increases MYC protein activity in the nucleus in a manner that is independent of protein abundance.

Similar results were obtained for *KDM6A*, a gene that encodes the H3K27me3 demethylase UTX, which cooperates with transcription and signaling factors to control gene expression programs^41,42^. As UTX is expressed in MCF7 cells (Fig. S1E), we knocked down endogenous *KDM6A* and re-expressed HA-tagged UTX protein, followed by RNA-seq analysis. To assess 3′UTR-dependent UTX protein activity, we investigated the genes that significantly increased or decreased upon *KDM6A* knockdown and where re-expression of KDM6A-NU or KDM6A-U reversed at least partially the knockdown effect. We detected nearly twice as many genes whose expression was at least partially rescued in the knockdown, when UTX protein was re-expressed from the U construct compared to its expression from the NU construct (Fig. 1K, Table S3). These results indicate that including the *KDM6A* 3′UTR in the mRNA template increases UTX protein activity in the nucleus in an abundance-independent manner. Taken together, these data show that the two tested HC 3′UTRs, when included in mRNA templates, influence the cellular activity of the encoded IDR-containing proteins in a protein abundance-independent manner. These results indicate an unexpected functional role for protein activity regulation by including HC 3′UTRs in expression vectors of two candidates.

### Including NC 3′UTRs in mRNA templates does not alter protein activity

To investigate if inclusion of NC 3′UTRs in expression vectors also affects activity of the mRNA-encoded proteins, we tested two enzymes. The *ENO1* gene encodes a glycolytic enzyme called α-enolase that increases lactate levels in the cell culture medium^43^. We knocked down endogenous *ENO1* in HeLa cells and measured the ability of NU and U constructs to rescue lactate production. We did not observe a difference in lactate accumulation between the two constructs (Fig. S1F-K). In addition, we tested protein activity of deoxyhypusine synthase (DHPS). The translation factor EIF5A contains a unique post-translational modification, called hypusination. DHPS performs the first step of hypusination, and its activity correlates with EIF5A hypusination level^44^. We knocked down endogenous *DHPS* in HeLa cells and transfected NU and U constructs of DHPS. We did not observe a strong difference in EIF5A hypusination level (Fig. S1L-P). For the two tested proteins, which do not contain IDRs, inclusion of the NC 3′UTRs in mRNA templates did not alter protein activity.

### Generation of a fully active JMJD3 histone demethylase in cells requires the *KDM6B* 3′UTR

Given these results, we set out to study how HC 3′UTRs — which are not part of proteins — could regulate protein activity. We focused on the *KDM6B* gene, which encodes the JMJD3 protein and functions as H3K27me3 demethylase in the nucleus. We chose this candidate as the protein has enzymatic activity, which allows us to quantify the expression level of transfected HA-tagged protein and the corresponding cellular activity in one assay. JMJD3 protein is expressed at very low levels in HeLa cells^45,46^. Therefore, we ectopically expressed KDM6B-NU or -U constructs and measured the reduction in H3K27me3 levels compared with untransfected cells using immunofluorescence (IF) staining (Fig. 1L).

With this assay, we do not measure enzymatic activity directly but use the reduction in H3K27me3 signal as cellular read-out for JMJD3 protein activity. In parallel, we quantify protein abundance of transfected HA-tagged JMJD3 using HA signal intensity (Fig. 1L). Cells transfected with NU or U constructs had comparable HA intensity values, suggesting similar protein levels, which was also observed when performing western blots (Fig. 1I, 1M). Despite no difference in JMJD3 protein abundance obtained from NU or U constructs, we observed a substantial difference in the reduction of H3K27me3 staining with the two constructs (Fig. 1N). Cells transfected with the CDS-only construct showed a modest reduction in methylation level with a median decrease of less than 10%. In contrast, cells transfected with the 3′UTR-containing vector showed a median decrease in H3K27me3 staining of 56% compared to untransfected cells (Fig. 1N). This result is consistent with substantially higher cellular JMJD3 activity, when the protein was translated from the 3′UTR-containing mRNA template.

### The endogenous *KDM6B* 3′UTR affects mesendoderm differentiation

Changes in H3K27me3 level are especially important during stem cell differentiation and embryonic development^45^. To rule out vector-dependent overexpression artifacts, we investigated whether the endogenous *KDM6B* 3′UTR is biologically important. We used CRISPR-Cas9 and a pair of guide RNAs to delete 875 base pairs of genomic sequence in human induced pluripotent stem cells (iPSCs). At the mRNA level, this deletion removes 68% of the *KDM6B* 3′UTR sequence without disrupting the CDS or mRNA 3′ end processing elements (Fig. S2A-C). The *Kdm6b* gene knockout in mouse embryonic stem cells, which fully prevents Jmjd3 protein expression, impairs mesoderm and subsequent cardiomyocyte differentiation^47^. We investigated whether partial deletion of the *KDM6B* 3′UTR would also result in phenotypic changes during iPSC to mesoderm differentiation. In the first three days of cardiomyocyte differentiation in culture, stem cells adopt mesoderm and mesendoderm markers, such as Brachyury, EOMES, and GATA6^47,48^. Partial deletion of the *KDM6B* 3′UTR (Udel) did not change endogenous JMJD3 protein levels, measured by western blot and flow cytometry (Fig. S2D, S2E). However, protein abundance of the three tested mesoderm/mesendoderm markers was significantly different in three homozygous Udel clones compared with three clones containing the wildtype 3′UTR (Fig. S2F-I). This result suggests that the endogenous *KDM6B* 3′UTR is important for faithful mesoderm and mesendoderm differentiation. The strongest 3′UTR-dependent difference was observed for GATA6 expression (Fig. S2H, S2I), which is one of the master regulators of heart development^48^, suggesting that the *KDM6B* 3′UTR could play a role during cardiomyocyte differentiation.

### The *KDM6B* 3′UTR is not required for activity regulation of the IDR-deleted JMJD3 protein

To examine the mechanism by which the *KDM6B* 3′UTR influences JMJD3 protein activity in a protein abundance-independent manner, we used the vector constructs. Full-length (FL) JMJD3 contains a long N-terminal IDR of over 1,100 residues and a structured C-terminus harboring a JmjC domain with demethylase activity^45,49^. To date, nearly all studies on JMJD3 activity have used a truncated construct that excludes the long N-terminal IDR and the *KDM6B* 3′UTR^45,49–52^. We call the IDR-deleted variant IDRdel (Fig. 2A, 2B).

**Figure 2.**
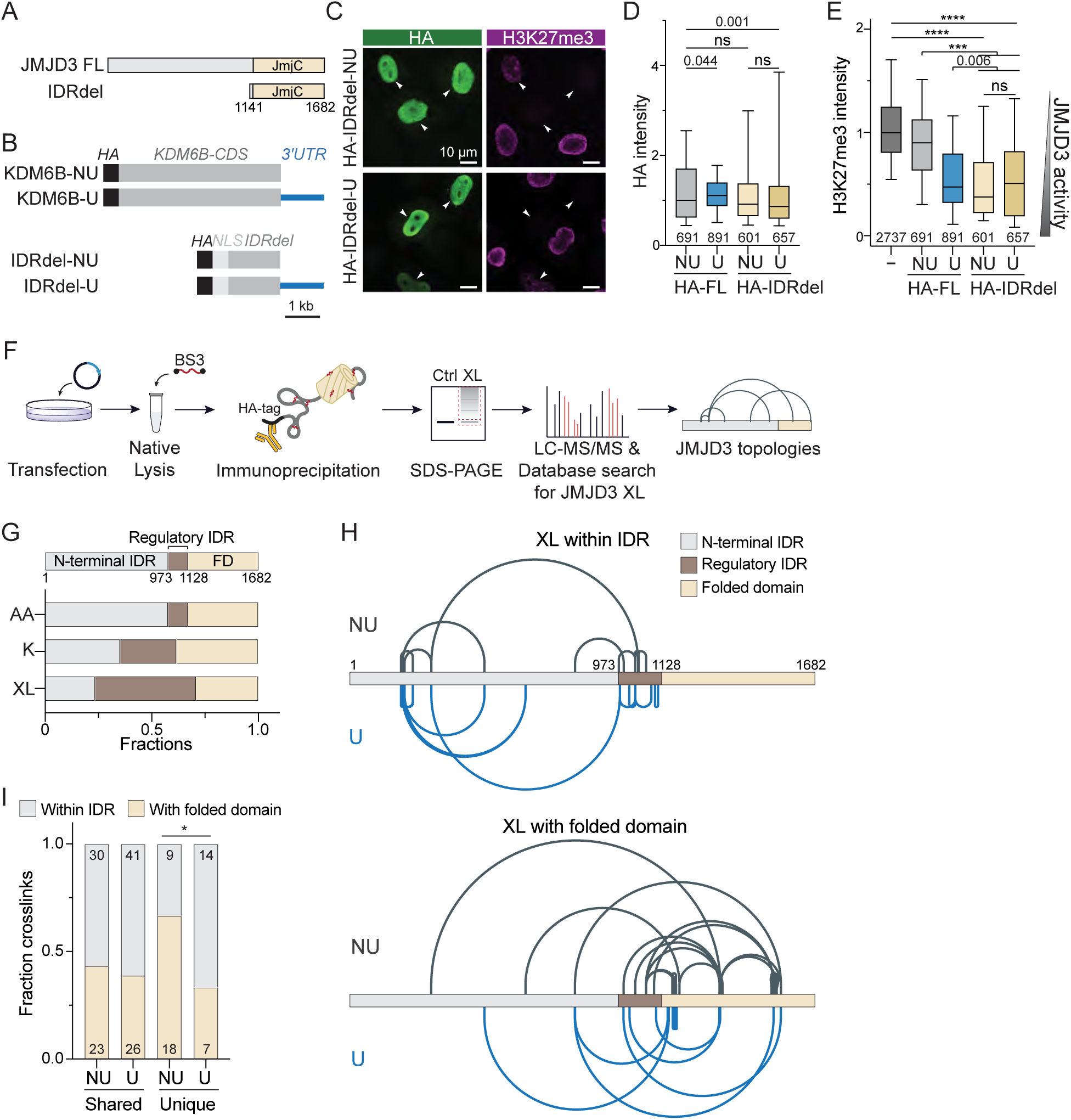
The 3′UTR-dependent difference in JMJD3 activity and protein conformation involve its IDR. **A.** Protein models for full-length (FL) and IDR-deleted (IDRdel) JMJD3. The structured JmjC domain is indicated in light brown, whereas the IDR is shown in gray. The amino acid boundaries of the IDRdel construct are shown. **B.** JMJD3 full-length and IDR deletion constructs are shown as in Fig. 1H. The SV40 nuclear localization signal (NLS) was added to the IDR deletion constructs. **C.** As in Fig. 1L, but 3′UTR-dependent activity was measured for the IDRdel protein using constructs from (B). **D.** As in Fig. 1M, but IDRdel protein abundance was measured as HA intensity from *N* = 3 independent experiments shown in (C), compared side-by-side with FL proteins. Mann-Whitney test: ns, not significant. **E.** As in Fig. 1N but shown are H3K27me3 values for FL and IDRdel-transfected cells from *N* = 3 independent experiments, as shown in (C). Mann-Whitney test: ns, not significant; ***, *P* = 7.8 x 10^−80^; ****, *P* < 10^−139^. **F.** Workflow for crosslinking mass spectrometry (XL-MS) to analyze JMJD3 conformation in cells. **G.** Distribution of all crosslinks (XL) that map to the indicated JMJD3 regions. The fraction of amino acids (AA), lysines (K), and XL in each region are shown. **H.** JMJD3 crosslinks uniquely detected after expression from NU (gray) or U (blue) construct are shown as lines connecting the crosslinked residues. Crosslinks within the IDR and crosslinks with the folded domain are plotted separately. **I.** Quantification of unique crosslinks shown in (H). Also shown are shared crosslinks (detected in both NU and U samples). Test for independent proportions, *P* = 0.02.

Repeating the IF assay with the IDR deletion constructs showed that protein expression of the full-length constructs and the IDR deletion constructs was similar, as determined by HA intensity (Fig. 2C, 2D). Expression of only the folded domain generates an active JMJD3 protein, resulting in a median reduction of 50% in H3K27me3 staining, which is comparable to the reduction observed with the U construct (Fig. 2E). Moreover, biogenesis of an active protein that only contains the folded domain does not require the *KDM6B* 3′UTR in the mRNA template (Fig. 2C-E). This experiment demonstrates a functional link between the JMJD3 IDR domain of the protein and the *KDM6B* 3′UTR in the mRNA: when both components are present or when both components are absent, a fully active protein is generated. Without the 3′UTR in the mRNA, the IDR negatively affects JMJD3 protein activity. In the native mRNA however, the repressive effect of the IDR is overcome by the 3′UTR.

### The *KDM6B* 3′UTR does not regulate JMJD3 protein localization to chromatin

Protein activity can be regulated by protein localization^6–8^. As we did not measure JMJD3 enzymatic activity directly, it is possible that the *KDM6B* mRNA 3′UTR controls JMJD3 subnuclear localization, and therefore its activity. As JMJD3 localizes to chromatin, we expressed our HA-tagged KDM6B constructs and performed Chromatin immunoprecipitation and sequencing (ChIP-seq) as high-resolution read-out for JMJD3 protein localization. The HA signal indicating JMJD3 localization to chromatin revealed reduced chromatin occupancy of the IDR-deleted compared with the full-length protein (Fig. S3A, S3B). However, importantly, the *KDM6B* 3′UTR had no effect on subnuclear localization of full-length JMJD3 protein (Fig. S3A, S3B), showing that JMJD3 protein localization is not regulated by the *KDM6B* 3′UTR.

In parallel, we performed H3K27me3 ChIP-seq to investigate the effects of IDR and mRNA 3′UTR on genome-wide cellular activity of JMJD3 (Fig. S3C, S3D). The ChIP-seq results confirmed the previous IF results, as we detected a strong reduction in H3K27me3 signal, consistent with full cellular JMJD3 activity in the IDR deletion construct or upon expression of the U construct (Fig. S3C, S3D). In contrast, the full-length protein generated from the NU vector only reduced the H3K27me3 signal by 50%, consistent with partial activity (Fig. S3C, S3D). This finding indicates that the *KDM6B* 3′UTR influences cellular JMJD3 protein activity in a manner that is independent of protein localization.

Protein activity can also be regulated by protein interactors, post-translational modifications or directly through protein conformation^53^. We performed SILAC-based immunoprecipitation followed by mass spectrometry of JMJD3 protein expressed from NU or U constructs and did not observe major differences in 3′UTR-dependent protein interactors or modifications (data not shown). Therefore, we hypothesized that the *KDM6B* 3′UTR alters JMJD3 protein conformation directly.

### The *KDM6B* 3′UTR in the mRNA template changes JMJD3 protein conformation

To test this hypothesis, we used NU and U constructs to express full-length JMJD3 protein in cells, followed by BS3 (bis(sulfosuccinimidyl)suberate)-mediated crosslinking mass spectrometry (XL-MS, Fig. 2F). To test potential 3′UTR-dependent changes in protein conformation, we focused our analysis on intra-JMJD3 crosslinks. As quality control, we examined crosslinking efficiency across samples by comparing the frequencies of looplinks, monolinks, and crosslinks. We did not observe significant differences between NU and U samples (Table S4), consistent with similar crosslinking efficiency.

We used all obtained crosslinks and determined their distribution across the JMJD3 IDR and folded domain. We observed that most IDR crosslinks mapped to a small region within the IDR, located upstream of the C-terminal folded domain. We call this region the putative ‘regulatory IDR’ as it only covers 9% of IDR sequence but contains 47% of IDR crosslinks (Fig. 2G).

Next, we focused on the unique crosslinks detected in the NU and U samples and observed a significant, 3′UTR-dependent difference in crosslinking pattern (Fig. 2H, 2I, Table S4). When JMJD3 protein was generated from an mRNA template containing the 3′UTR, two-thirds of unique crosslinks were indicative of IDR-IDR interactions, whereas in the absence of the 3′UTR two-thirds of unique crosslinks involved binding to the folded domain that contains the enzymatic activity (Fig. 2I). This experiment provides direct evidence for a 3′UTR-dependent difference in JMJD3 protein conformation involving its IDR. It suggests that the presence of the *KDM6B* 3′UTR promotes interactions within the IDR, whereas its absence induces protein folding across domains, that is, between the IDR and the folded domain.

### The hydrophobic region in the regulatory IDR has a repressive effect on JMJD3 protein activity

To learn how the 3′UTR in the mRNA template could change IDR-dependent protein conformations, we set out to learn the regulatory logic of the IDR. We examined the amino acid composition of the IDR regulatory domain and observed that it contains a highly positively charged stretch with a net charge of +16 and an adjacent region with a high fraction of hydrophobic (HP) amino acids (Fig. 3A, S3E). Moreover, PhyloP sequence conservation analysis revealed that the HP region has the highest sequence conservation within the entire IDR and its sequence is largely conserved across vertebrate species, including zebrafish (Fig. 3A).

**Figure 3.**
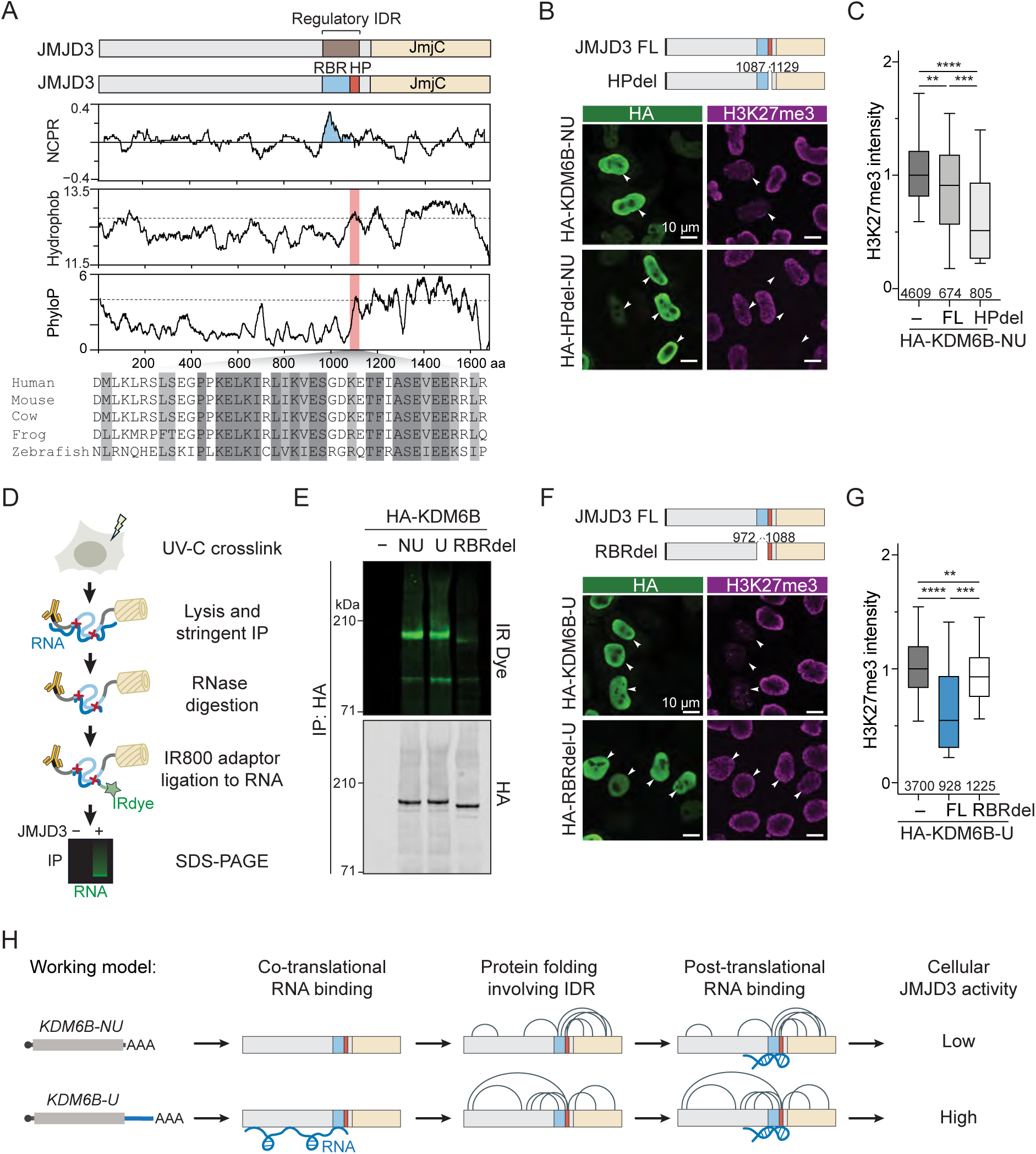
The JMJD3 IDR contains regions with opposite effects on JMJD3 protein activity. **A.** The regulatory IDR consists of a putative RNA-binding region (RBR, aa 973-1087, blue) and a conserved hydrophobic region (HP, aa 1088-1128, red). Top: net charge per residue (NCPR). Middle: hydrophobic scale values per residue. Bottom: 100-way PhyloP scores of KDM6B CDS. Amino acid alignment of HP region across selected species are shown below. Dark gray, perfect conservation; light gray, conservative substitution. **B.** As in Fig. 1L, but JMJD3 abundance and cellular activity assay of full-length and HPdel is shown. **C.** As in Fig. 1N, but normalized H3K27me3 intensity values obtained from *N* = 3 independent experiments are shown for constructs from (B). Mann-Whitney test: **, *P =* 2.8 x 10^−17^; ***, *P* = 5.0 x 10^−23^; ****, *P =* 3.1 x 10^−133^. **D.** irCLIP workflow for the detection of RNA-JMJD3 protein interaction in cells. **E.** irCLIP of the indicated HA-tagged KDM6B constructs expressed in HeLa cells. **F.** As in Fig. 1L, but JMJD3 abundance and cellular activity assay of full-length and RBRdel is shown. **G.** As in Fig. 1N, but the cellular activity of full-length JMJD3 and RBRdel was measured as reduction in H3K27me3 from *N* = 3 independent experiments. Mann-Whitney test: **, *P =* 5.5 x 10^−17^; ***, *P* = 2.6 x 10^−77^; ****, *P =* 1.4 x 10^−137^. **H.** Working model for differential effect of co- and post-translational RNA-IDR interaction in the regulation of 3′UTR-dependent cellular JMJD3 activity. Details, see text.

We deleted the HP region and tested JMJD3 protein activity by IF and observed a strong decrease in H3K27me3 staining, even when expressed from an mRNA template lacking the 3′UTR (Fig. 3B, 3C, S3F). This result suggests that the HP region in the IDR has a strong repressive effect on cellular JMJD3 activity, when the protein is translated in the absence of the mRNA 3′UTR.

### The IDR RNA-binding region is necessary for 3′UTR-mediated regulation of JMJD3 protein activity

It was previously shown that positively charged regions in IDRs can act as RNA-binding regions (RBR)^33,34^. To examine whether the JMJD3 IDR region with a high positive net charge acts as RBR in cells, we deleted the putative RBR and performed irCLIP^54,55^. We transfected our constructs, performed RNA-protein crosslinking in cells, followed by HA immunoprecipitation and visualization of the crosslinked RNA by an RNA-ligated dye (Fig. 3A, 3D). Deletion of the RBR strongly reduced the RNA signal, when compared with the full-length JMJD3 protein (Fig. 3E). This experiment shows that this IDR region is the major RNA-binding region in JMJD3 protein.

To test whether the RBR is important for cellular JMJD3 protein activity, we expressed the RBR deletion construct from a 3′UTR-containing mRNA template and performed IF (Fig. 3F). Whereas the full-length JMJD3 protein generated from the U construct strongly reduced H3K27me3 staining, absence of the RBR showed only weak reduction (Fig. 3F, 3G, S3G). This result shows that the RBR is necessary for cellular JMJD3 activity, when the protein was generated from a 3′UTR-containing mRNA template and suggests that the RBR in the JMJD3 IDR mediates the effect of the mRNA 3′UTR.

So far, we learned that the RNA-binding region in the JMJD3 IDR is necessary for 3′UTR-dependent control of cellular JMJD3 activity (Fig. 3G). However, this region is not sufficient, as both versions of full-length JMJD3 protein — generated from the NU or U constructs — contain this region, but only the protein generated from the U construct shows full activity. As both protein versions bind to RNA in cells, we propose that there is a difference between post-translational and co-translational RNA binding. In our working model, only presence of the mRNA 3′UTR allows co-translational binding of the 3′UTR and the IDR in the JMJD3 nascent chain, thus being capable of directly changing its protein conformation (Fig. 3H).

### RNA-IDR binding changes the interaction preference between the JMJD3 IDR and the folded domain in vitro

To examine if RNA can directly change JMJD3 protein conformation, we used recombinant JMJD3 protein domains and assessed RNA-dependent interactions. We focused on the regulatory IDR and the folded domain, as these domains are predominantly involved in intra-JMJD3 contacts according to the XL-MS experiment (Fig. 2H, 2I). We used in vitro protein crosslinking to test the influence of in vitro-transcribed RNA on IDR-IDR and IDR-folded domain interactions.

We generated the regulatory IDR domain and the C-terminal folded domain as separate protein domains (Fig. 4A, S4A). First, we used fluorescence polarization assays to test in vitro RNA binding of the two domains. At the tested concentrations, we observed no RNA binding to the folded domain, but strong binding to the regulatory IDR at physiological salt concentration (Fig. S4B-D). Deletion of the RBR in the IDR strongly reduced RNA binding (Fig. S4E). The RBR has a very high positive net charge but lacks known RNA-binding motifs (Fig. S4F). Moreover, different in vitro transcribed RNAs interacted with the IDR with similar affinity, suggesting that the RNA-IDR interactions are not sequence specific but rather mediated by electrostatic interactions (Fig. S4B-E).

**Figure 4.**
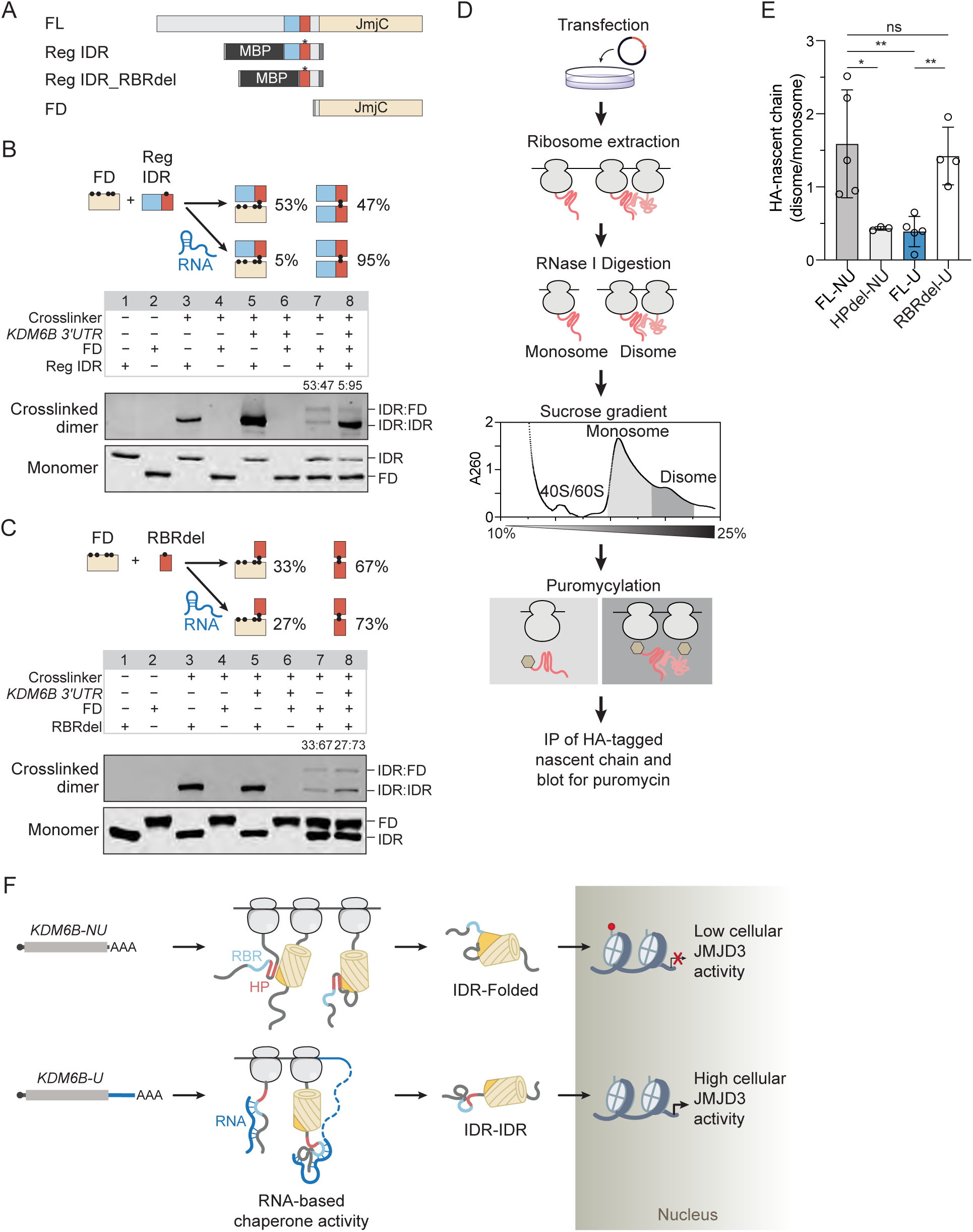
RNA-IDR interaction is necessary and sufficient for a change in JMJD3 domain interaction preference. **A.** JMJD3 protein domain constructs used for RNA-binding and thiol-thiol crosslinking experiments in vitro. All constructs contain an N-terminal 6xHis-tag (medium gray) and C-terminal Strep-tag (dark gray). The three native cysteines in the regulatory IDR were mutated, while a single cysteine was introduced at E1102 (asterisk) to enable crosslinking. **B.** Immunoblots showing RNA-dependent protein-protein interactions between recombinant JMJD3 regulatory IDR and folded domain (FD) detected by BMOE-mediated thiol-thiol crosslinking. To resolve different dimer bands, a 6% Tris-Glycine gel was used, which does not capture the monomer bands. The same reactions were run on a separate 8% Tris-Glycine gel to resolve the monomer bands. An 8% gel showing monomers and dimers on one gel is presented in Fig. S4G. **C.** As in (B) but using a construct without the RNA-binding region (RBRdel) instead of the whole regulatory IDR to detect RNA-dependent protein-protein interactions. The reaction was run on an 8% Tris-Glycine gel. The full gel is presented in Fig. S4H. **D.** Workflow of co-translational disome assay. HeLa cells transfected with the indicated constructs were lysed, treated with ribonuclease, and separated into monosome and disome fractions using sucrose gradient centrifugation. Nascent chains in each fraction were labeled with puromycin (yellow hexagon). HA-tagged target protein was immunoprecipitated, newly synthesized target was detected by immunoblotting with anti-puromycin, and results were normalized to RPLP0. **E.** Abundance of HA-tagged nascent chains after transfection of the indicated constructs in disome over monosome fractions. Puromycylated HA-tagged JMJD3 band intensity in immunoprecipitated fractions was normalized to RPLP0 band intensity in the input to account for different ribosome amounts in each fraction. Shown are mean ±LSD from *N* = 5 independent experiments. Raw data are shown in Fig. S4J. Unpaired t-test: ns, not significant; *, *P* < 0.05; **, *P* < 0.01. **F.** Model for 3′UTR-dependent regulation of JMJD3 cellular activity through differences in protein folding accomplished by co-translational RNA-mediated chaperone activity provided by the *KDM6B* 3′UTR. Details, see text.

To test RNA-dependent changes in protein domain interactions, we performed a bismaleimidoethane (BMOE)-mediated thiol-thiol crosslinking assay (Fig. 4B). We used native cysteines in the folded domain and mutated all three naturally occurring cysteines in the regulatory IDR outside of the HP region and introduced a single cysteine within the HP region at E1102C (Fig. 4A). The folded domain alone did not form crosslinked product and addition of different RNAs (*KDM6B or HSPA1B* 3′UTRs) had no effect, consistent with its inability to bind RNA (Fig. 4B, lanes 4 and 6, S4G).

In the absence of RNA, mixing of folded domain and regulatory IDR generated near equal amounts of IDR-IDR and IDR-FD dimers (Fig. 4B, lane 7, S4G). Addition of RNA to the mixture generated a 95:5 dimer ratio, by promoting the IDR-IDR interaction (Fig. 4B, lane 8, S4G). This experiment shows that the presence of RNA promotes IDR-IDR interaction, whereas absence of the RNA allows formation of equal amounts of IDR-IDR dimers and IDR-folded domain dimers (Fig. 4B). The interaction pattern observed with recombinant protein domains perfectly mimics the domain interactions observed by XL-MS obtained from cells, where the full-length proteins were generated from mRNA templates containing or lacking the *KDM6B* 3′UTR (Fig. 2H, 2I).

In contrast, performing the in vitro crosslinking assay in the absence of the RBR in the IDR abrogated the RNA-mediated increase in IDR-IDR interaction (Fig. 4C, S4H), indicating that both the RNA and the RNA-binding region need to be present simultaneously for the switch in domain interaction preference towards IDR-IDR interaction. This experiment demonstrates that RNA-IDR binding is necessary and sufficient for the change in domain interaction preference between the JMJD3 IDR and its folded domain in vitro. As the in vitro crosslinking experiment recapitulated the 3′UTR-dependent increase in IDR-IDR interaction at the expense of IDR-folded domain interaction observed by XL-MS, performed on cellular full-length JMJD3 protein, these results suggest that the *KDM6B* 3′UTR could act as chaperone for the JMJD3 IDR and directly change JMJD3 conformations co-translationally.

### Simultaneous presence of the *KDM6B* 3′UTR and the IDR RNA-binding region are required for co-translational effects on JMJD3 nascent chain interactions

Next, we tested whether the *KDM6B* 3′UTR can alter cellular JMJD3 protein conformation in a co-translational manner. The XL-MS results on cellular JMJD3 (Fig. 2H, 2I) cannot distinguish whether the interactions occur between molecules or within a single protein molecule. AlphaFold3 predicts that JMJD3 forms a domain-swapped dimer, where the HP region binds to the folded domain of another JMJD3 molecule (Fig. S4I)^56,57^. Domain-swapped dimers often are generated through co-translational protein complex assembly^58–60^.

Therefore, we developed a ‘co-translational disome assay’ to investigate a potential 3′UTR-dependent change in co-translational JMJD3 protein conformation (Fig. 4D). Disomes are pairs of ribosomes and can occur pathologically, through stress-associated ribosome collisions, or physiologically, through nascent chain interaction from neighboring ribosomes^59,61,62^. After transfection of our constructs, we used sucrose gradient centrifugation to extract monosomes and disomes from cells, used puromycin to label the nascent chains in each fraction, and immunoprecipitated the JMJD3 nascent chains with HA antibody, followed by normalization to RPLP0, a ribosomal protein, to adjust for differences in input in the isolated fractions (Fig. S4J). We ruled out that the disomes are generated through ribosome collisions, as we did not observe a difference in ZAK phosphorylation, which correlates with stress-associated ribosome collisions (Fig. S4K, S4L)^61^.

Presence of HA-tagged JMJD3 in disomes indicates co-translational homodimerization, whereas its presence in monosomes indicates monomeric protein^59,60^. The HA signal of the different constructs was plotted as disome-over-monosome ratio. With the full-length protein obtained from the NU construct, we observed significantly more JMJD3 nascent chains in disomes compared with monosomes (Fig. 4E). By contrast, expression of the full-length protein from the U construct shifted the majority of JMJD3 nascent chains to the monosome fraction, indicative of decreased JMJD3 homodimerization (Fig. 4E). These results show that the 3′UTR in the mRNA template has a co-translational effect on JMJD3 nascent chain interactions.

To test whether the regulatory IDR is involved in co-translational changes in nascent chain interactions, we examined the HP and RBR deletion constructs. Whereas deletion of the HP region decreased interactions, deletion of the RBR increased co-translational protein-protein interactions (Fig. 4E). These results show that both the 3′UTR in the mRNA and the RNA-binding region in the IDR need to be present to avoid the co-translational nascent chain interactions mediated by the HP region in the NU translated protein (Fig. 4E).

These results are consistent with a model (Fig. 4F), in which the 3′UTR co-translationally binds to the IDR and directly changes domain interaction preference. As the 3′UTR results in a change in cellular protein activity in the nucleus, it seems that the HP region in the IDR could interfere with folding of the structured domain, thus, resulting in a permanent change in JMJD3 protein conformation that is determined during translation. Taken together, our results suggest that the *KDM6B* 3′UTR has RNA-based IDR chaperone activity during JMJD3 protein folding.

### Not all 3′UTRs function as RNA-mediated chaperones for JMJD3

Our in vitro RNA-protein binding assay suggests that the interaction between RNA and the JMJD3 IDR is mediated by electrostatic interactions (Fig. S4D, S4E). Moreover, the in vitro crosslinking assays suggests that the RNA-dependent change in protein domain interaction preference is independent of RNA sequence (Fig. S4G, S4H). Based on these observations, we investigated whether other 3′UTRs, when fused to the *KDM6B* coding sequence, can function as RNA-based JMJD3 IDR chaperones in cells. In the vector constructs, we swapped the *KDM6B* 3′UTR with four different 3′UTRs and assessed cellular JMJD3 activity through a reduction of H3K27me3 staining by IF (Fig. 5A). We tested 3′UTRs that are not highly conserved (Table S1). For two of the tested 3′UTRs (derived from *IDH1* or *HSPA1B* mRNA), we observed substantially reduced H3K27me3 staining, consistent with these 3′UTRs containing IDR chaperone activity. However, the other two 3′UTRs (derived from *PIN1* or *MAP2K2* mRNA) behaved like the NU construct and reduced H3K27me3 staining only to a small extent, suggesting that these 3′UTRs do not function as IDR chaperones (Fig. 5A, 5B, S5A, S5B).

**Figure 5.**
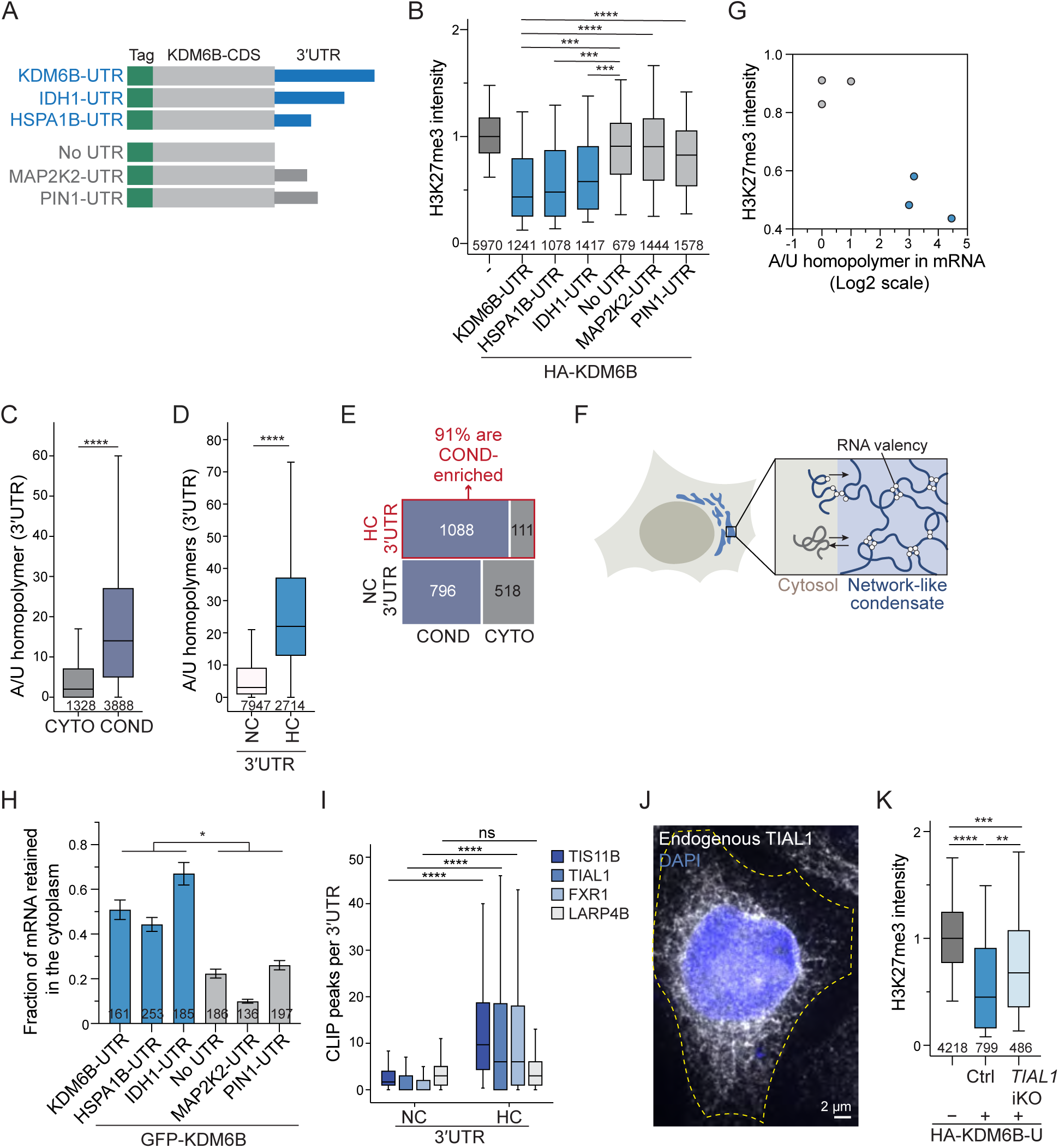
3′UTRs with RNA-based chaperone activity are multivalent and enriched in meshlike condensates. **A.** GFP- or HA-tagged KDM6B constructs for 3′UTR-swapping experiments. The KDM6B CDS was fused with four different 3′UTRs. 3′UTRs with IDR chaperone activity are colored blue, whereas 3′UTRs without IDR chaperone activity are colored gray. **B.** As in Fig. 1N, but 3′UTR-dependent JMJD3 cellular activity was measured for the constructs shown in (A) from *N* = 3 independent experiments. A reduction in H3K27me3 intensity indicates active JMJD3. Mann-Whitney test, ***, *P* < 10^-39^; ****, *P* < 10^-84^. **C.** RNA valency (number of A/U homopolymers) in the 3′UTRs of mRNAs enriched in the cytosol (cyto, *N* = 1328) or in meshlike condensates (cond, *N* = 3888). Mann-Whitney test, 8.4 x 10^-182^. **D.** As in (C) but shown for HC 3′UTRs (*N* = 2714) and NC 3′UTRs (*N* = 7947). Mann-Whitney, *P* = 0. **E.** Contingency table showing the relationship between mRNAs containing HC and NC 3′UTRs and their subcytoplasmic localization to cytosol (cyto) or meshlike condensates (cond). X^2^ = 302.38, *P* < 0.0001. **F.** Schematic of a cell with cytoplasmic meshlike condensate (blue). Zoom-in depicts entrapment of multivalent mRNAs in meshlike condensates, which are generated through RNA-RNA interactions of multivalent mRNAs. In contrast, low-valency mRNAs are not entrapped in meshlike condensates, thus mostly localizing to the cytosol. **G.** Relationship between 3′UTR multivalency as indicated by the number of A/U homopolymers and 3′UTR-dependent cellular JMJD3 activity, measured by reduction in H3K27me3 level, obtained from (B). **H.** Digitonin RNA-FISH results obtained from the indicated transfected GFP-tagged constructs from (A) in HeLa cells. The fraction of RNA-FISH signal obtained after digitonin treatment is shown as mean ± SEM from *N* = 3 independent experiments. The number of analyzed cells is shown. T-test for independent samples, *, *P* = 0.014. **I.** CLIP peaks of the indicated condensate-enriched RNA-binding proteins detected for NC and HC 3′UTRs. LARP4B is shown as negative control, as it is not condensate enriched. Mann-Whitney test, ****, *P* < 10^−243^; ns, not significant. **J.** Immunofluorescence staining of endogenous TIAL1 protein in HeLa cells. DAPI was used to stain the nucleus. The yellow dotted line indicates the cell boundary. A representative image is shown. **K.** As in Fig. 1N, but 3′UTR-dependent JMJD3 cellular activity was measured in doxycycline-induced *TIAL1* knockout (iKO) versus control (Ctrl) HeLa cells containing non-targeting guide RNAs. *N* = 3 independent experiments were performed. A reduction in H3K27me3 intensity indicates active JMJD3. **, *P =* 1.4 × 10^−13^; ***, *P* = 5.7 × 10^−40^; ****, *P =* 3.0 × 10^−130^.

### Highly conserved 3′UTRs are multivalent and predicted to localize to meshlike condensates

To identify RNA features that distinguish 3′UTRs with or without IDR chaperone activity, we compared the RNA features between HC and NC 3′UTRs and found that they substantially differ with respect to enriched 6-mer motifs, 3′UTR length, AU content, mean free energy of their predicted secondary structures, and RNA-RNA binding site density (Fig. S5C-H, Table S1)^4^. Several of these RNA features correlate with each other and have been associated with multivalent RNAs (Table S5)^12^.

RNA valency is a parameter that describes accessible RNA-RNA interaction sites and multivalent RNAs were previously shown to be sufficient to generate meshlike condensates in vitro^12^. RNA valency is determined by both RNA length and structure^12,63,64^. We searched for a single parameter that could serve as surrogate marker for multivalent cellular 3′UTRs, which differ substantially in length. We observed that the sum of A or U homopolymers with at least five consecutive nucleotides (also called A5/U5) in a given RNA correlates strongly with both

RNA length and weak predicted RNA structure and therefore, combines the two essential features of multivalent RNAs (Table S5). Moreover, as this parameter separates well the two groups of RNAs that were initially used to define RNA valency experimentally (Fig. S5I)^12^, we are using the number of A5/U5 in a given 3′UTR as surrogate marker for 3′UTR valency. A5/U5 distinguishes well between cytosolic and meshlike condensate-enriched mRNAs obtained from cells (Fig. 5C)^13,14^ and separates nearly perfectly HC from NC 3′UTRs (Fig. 5D). Intersecting the group of HC 3′UTRs with meshlike condensate-enriched mRNAs revealed a very strong overlap, as 91% of HC 3′UTRs are condensate enriched (Fig. 5E). Taken together, our data suggest that HC 3′UTRs are multivalent, allowing the 3′UTR-containing mRNAs to become entrapped in meshlike condensates in cells (Fig. 5F). In contrast, mRNAs with NC 3′UTRs have low valency, which prevents entrapping in meshlike condensates and promotes their cytosolic localization^12^.

### The tested multivalent 3′UTRs act as JMJD3 IDR chaperones

Importantly, using A5/U5 as surrogate marker for RNA multivalency, distinguishes perfectly the RNAs with and without IDR chaperone activity used in the 3′UTR-swapping experiment. 3′UTRs that increased cellular JMJD3 protein activity had multivalent 3′UTRs, as they had 7, 8, or 21 A/U homopolymers in their 3′UTRs. In contrast, 3′UTRs that were unable to increase JMJD3 protein activity had low-valency 3′UTRs, as they only contained 0 or 1 A/U homopolymers (Fig. 5G).

To investigate whether multivalent 3′UTRs enable mRNA enrichment in meshlike condensates in cells, we performed a previously established ‘digitonin RNA-FISH assay’^13^. With this assay, endogenous cytosolic and meshlike condensate-enriched mRNAs can be distinguished experimentally. Digitonin permeabilizes the plasma membrane but leaves internal membranes intact^65^. Therefore, digitonin treatment extracts unbound cytosolic mRNAs, whereas meshlike condensate-enriched mRNAs are largely retained in the cytoplasm^13^.

Here, we performed this assay using mRNAs derived from transfected expression vectors. mRNA valency of 3′UTR-containing *MYC*, *KDM6A*, and *KDM6B* mRNAs is at least two-fold higher compared to their 3′UTR-deleted counterparts (Fig. S6A) and digitonin RNA-FISH showed that the 3′UTR-containing mRNAs are largely retained in the cytoplasm, whereas deletion of their multivalent 3′UTRs substantially decreased their cytoplasmic mRNA retention. This result is consistent with mRNA enrichment in meshlike condensates by HC 3′UTRs (Fig. S6B, S6C).

A similar pattern was observed for the 3′UTRs used in the 3′UTR-swapping experiment. The mRNAs derived from constructs that had multivalent 3′UTRs and acted as IDR chaperones (*IDH1* and *HSPA1B* 3′UTR) were significantly more retained in the cytoplasm upon digitonin treatment than the constructs with 3′UTRs that lacked IDR chaperone activity (Fig. 5H, S6D). Taken together, for eight tested 3′UTRs, we observed a strong correlation between 3′UTR multivalency, enrichment in cytoplasmic meshlike condensates, and RNA-based chaperone activity (Fig. 5C, 5D, 5G, 5H, S6A-D), suggesting that at least some condensate-enriched RNAs have RNA-mediated chaperone activity.

### RNA-mediated IDR chaperone activity for JMJD3 requires the condensate-enriched RNA-binding protein TIAL1

As meshlike condensates are generated by specific RNA-binding proteins together with their bound mRNAs^11–14^, we investigated whether condensate-enriched RNA-binding proteins, such as TIS11B, TIAL1, or FXR1 are necessary for RNA-mediated chaperone activity^13,14^. We examined the number of 3′UTR-mapped CLIP peaks of these RNA-binding proteins between HC and NC 3′UTRs and observed significantly more peaks, which correspond to binding sites of the RNA-binding proteins, in HC than in NC 3′UTRs (Fig. 5I). LARP4B served as negative control, as it is not enriched in meshlike condensates (Fig. 5I)^13^.

It was previously shown that the *KDM6B* 3′UTR is strongly bound by TIAL1 (Table S1)^66^. Moreover, IF staining of endogenous TIAL1 in HeLa cells shows a reticulated, perinuclear localization pattern, characteristic for its enrichment in meshlike condensates (Fig. 5J)^11–13^. We investigated whether TIAL1 is necessary for the *KDM6B* 3′UTR to act as IDR chaperone. We transfected the U construct in HeLa cells containing an inducible *TIAL1* knockout and observed less reduction in H3K27me3 levels in cells lacking TIAL1, compared with control cells (Fig. 5K). This result suggests lower IDR chaperone activity of the *KDM6B* 3′UTR in the absence of the condensate-enriched RNA-binding protein (Fig. 5K, S6E-G) and is consistent with a model by which the multivalent *KDM6B* 3′UTR acts as RNA-based IDR chaperone within meshlike condensates.

### Hydrophobic amino acid clusters in the JMJD3 IDR have folding potential

Lastly, we set out to obtain a better understanding of the JMJD3 IDR features that require the presence of RNA acting as IDR chaperone. The regulatory region in the JMJD3 IDR contains a strong basic patch, followed by a hydrophobic region with clustered HP residues (Fig. 6A). We searched for additional regions within the JMJD3 IDR that contain HP amino acid clusters adjacent to basic patches and observed two more regions (Fig. 6A). We intersected the combined IDR basic and HP clusters with the XL-MS data and obtained a striking result. Although these regions only represent 16% of IDR residues, they harbor 77% of IDR crosslinks (Fig. 6B), suggesting that these IDR regions have a high folding potential.

**Figure 6.**
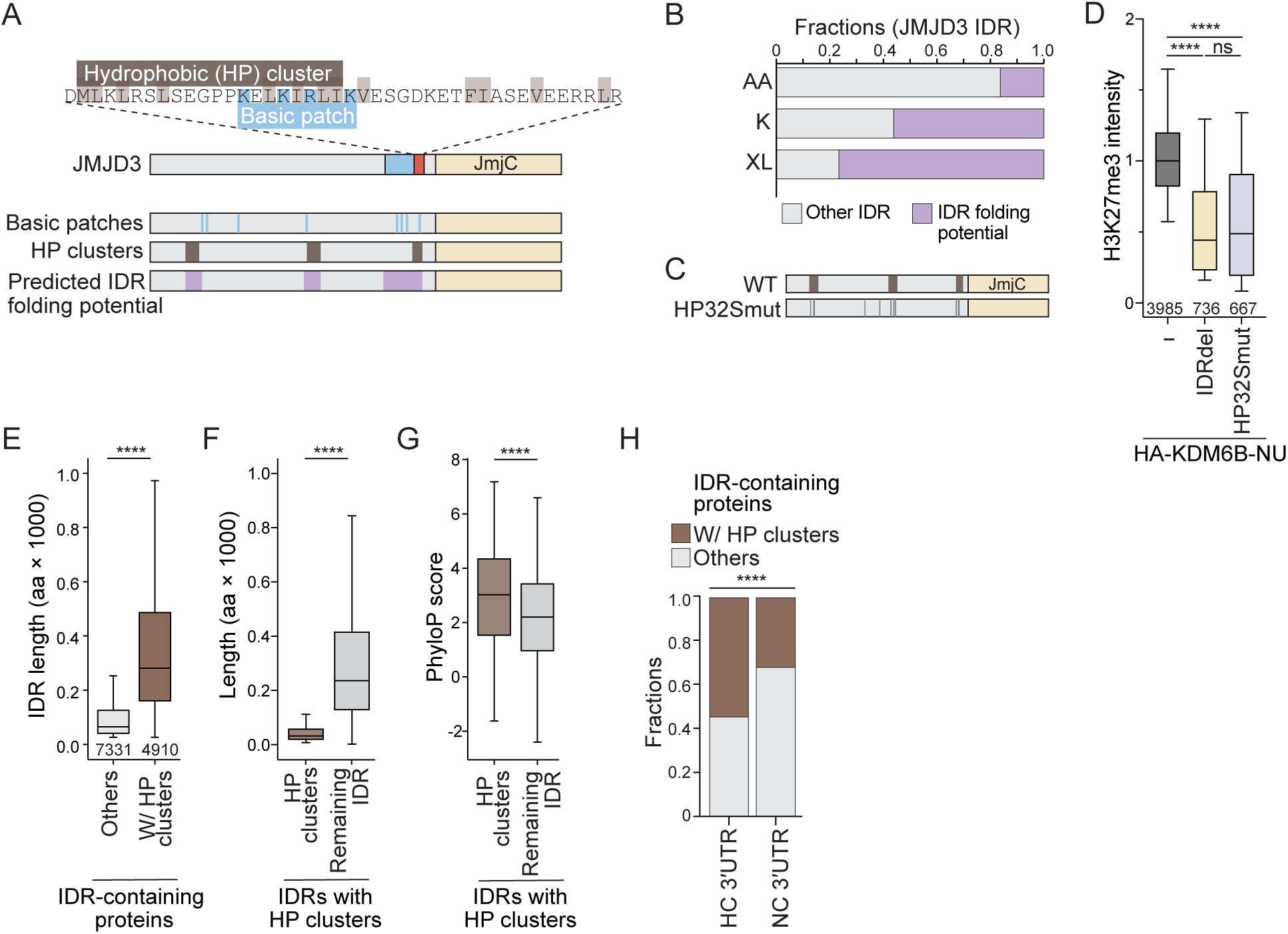
Hydrophobic amino acid clusters in the IDR participate in protein folding of JMJD3. **A.** Mapping of hydrophobic clusters and basic patches in the JMJD3 IDR. **B.** Distribution of crosslinks between regions in the JMJD3 IDR with predicted folding potential (purple) from (A) and the remaining IDR amino acids (AA). The distribution of lysine residues is also shown (K). **C.** HP32Smut construct contains mutation of 32 hydrophobic residues to serines (gray bars). The mutated residues map to hydrophobic amino acid clusters (brown blocks) in the JMJD3 IDR. **D.** As in Fig. 1N, but JMJD3 cellular activity was measured as reduction in H3K27me3 for IDRdel- or HP32Smut-transfected cells from *N* = 3 independent experiments. Mann-Whitney test: ****, *P* < 10^−131^. **E.** For each IDR-containing protein the longest IDR was analyzed (*N* = 12,241). Shown is IDR length distribution of IDRs with (brown; *N* = 4,910) or without HP clusters (gray; *N* = 7,331). Mann-Whitney test: ****, *P* =0. **F.** For IDRs with HP clusters (*N* = 4,910), amino acid length of the HP clusters and the remaining IDR region is shown. Wilcoxon test: ****, *P* = 0. **G.** As in (F), but 100-way PhyloP scores of CDS regions that encode the HP clusters versus the remaining IDR regions are shown. Wilcoxon test: *P* = 5.1 x 10^-315^. **H.** Fractions of mRNAs that encode proteins with HP clusters in their IDRs are shown for mRNAs encoding IDR-containing proteins, with HC (*N* = 2,269) or NC 3′UTRs (*N* = 4,279). Chi-Square test of independence, *X*^2^ = 310, ****, *P* < 0.00001.

This hypothesis is supported by the following experimental data. The JMJD3 IDR contains 210 hydrophobic residues, 32 of which are located within the HP clusters. Mutation of these residues to serines (HP32Smut) generated a protein that had equal cellular activity as JMJD3 lacking the entire IDR (Fig. 6C, 6D, S7A, S7B). Although the protein generated from the HP32Smut construct contains more than 1,100 IDR residues, it is fully active regarding cellular reduction of H3K27me3 levels, suggesting that the JMJD3 IDR without HP clusters does not have folding propensity (Fig. 6C, 6D). This experiment is consistent with a model in which the HP clusters in the JMJD3 IDR could interfere with folding of the structured domain during co-translational protein folding.

### Hydrophobic amino acid clusters in IDRs have high sequence conservation scores

To identify proteome-wide IDRs with HP clusters, we focused on the longest IDR of each protein (Table S1). We identified 4910 IDRs (corresponding to 40% of tested IDRs) with at least one HP cluster. HP clusters are characteristic of long IDRs with a median length of 281 amino acids (Fig. 6E). Within IDRs, HP clusters only occupy small regions, corresponding to 10-15% of IDR sequence (Fig. 6F). However, within each IDR, the sequences that encode HP clusters have significantly higher PhyloP conservation scores than the sequences encoding the remaining IDR residues, suggesting that HP clusters are functionally relevant (Fig. 6G). Lastly, mRNAs that encode proteins with HP cluster-containing IDRs are strongly enriched for HC 3′UTRs (Fig. 6H).

The molecular features of the accompanying IDR basic patches are currently less clear. Basic patches occur more frequently in IDRs than in folded domains and IDRs with HP clusters have more basic patches (Fig. S7C-E). However, it is currently unknown whether basic patches in IDRs require a minimum distance or minimum net charge to enable RNA-based IDR chaperone activity.

### 3′UTR-dependent changes in protein states occur in candidates with hydrophobic clusters in their IDRs

Taken together, we postulate that proteins that lack IDRs and proteins without HP clusters in their IDRs fold independently of mRNA 3′UTRs. In contrast, proteins with HP clusters in their IDRs potentially require the presence of RNA-mediated chaperones during translation. This implies that protein expression from vectors that include or lack their mRNA 3′UTRs could change protein folding in cells.

To test this prediction, we transfected CDS vectors with or without full-length 3′UTRs and performed in-cell crosslinking with Disuccinimidyl Glutarate (DSG), followed by western blot analysis (Fig. 7A). We tested four candidate mRNAs, whose encoded proteins do not contain IDRs (ENO1, DHPS, RACK1, EIF3I) and three candidate mRNAs, whose encoded proteins contain IDRs without HP clusters (SSBP4, SSRP1, SF3B4; Table S6). We performed western blots in the absence and presence of crosslinking reagent and used line profiles and Pearson correlation coefficients to quantify potential differences in protein crosslinking pattern (Fig. 7A-E, S7F, S7G). In all tested cases, the crosslinking patterns were nearly identical, when the proteins were expressed from CDS vectors containing or lacking their corresponding 3′UTRs (Fig. 7B, 7D, 7E, S7F, S7G). These results suggest that for these candidates, the mRNA 3′UTRs do not influence the observed protein crosslinking pattern, which is a low-resolution read-out for potential differences in protein folding, oligomerization states, or protein interaction partners. All the tested candidates have low-valency NC 3′UTRs (Table S6).

**Figure 7.**
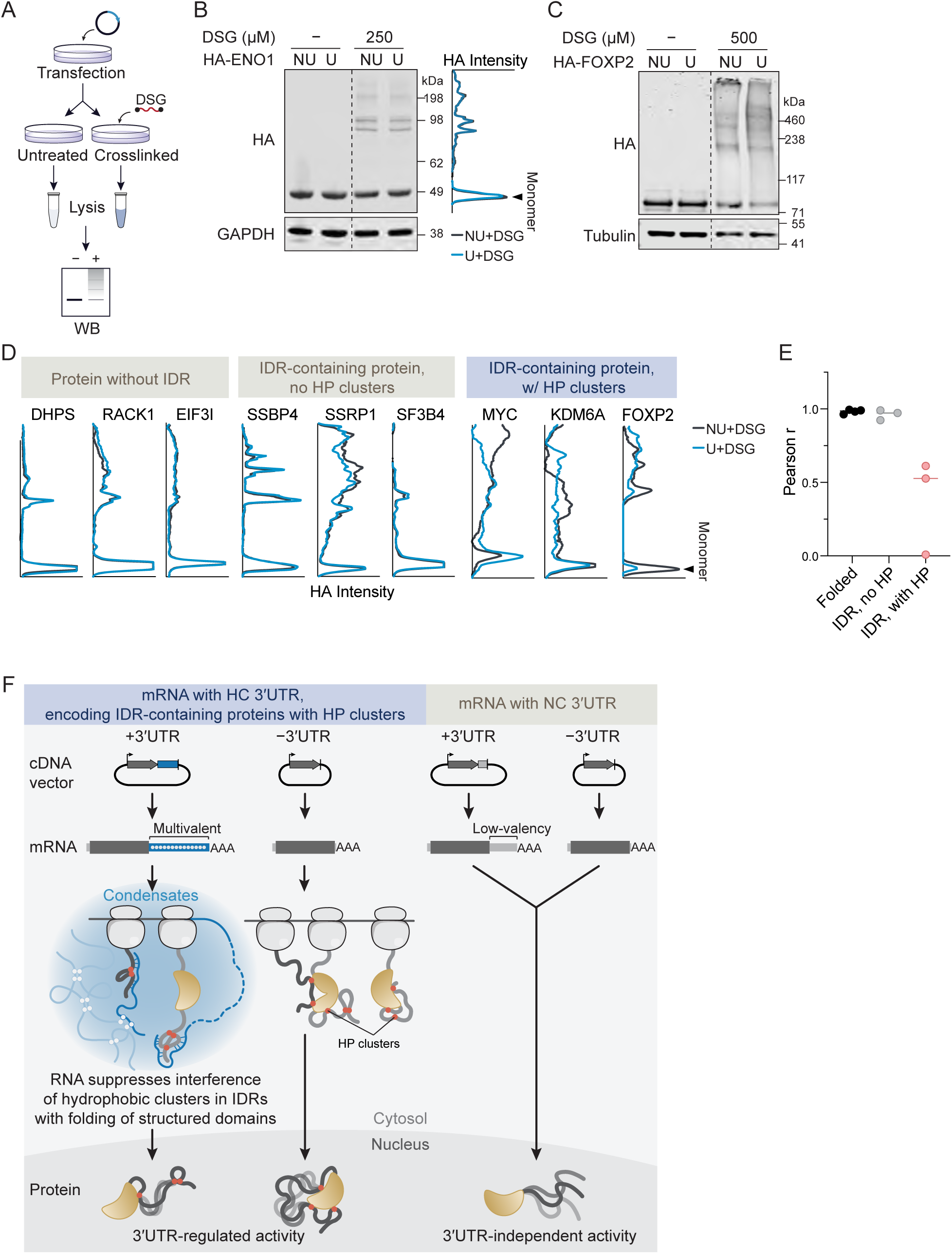
The protein states of proteins with hydrophobic clusters in their IDRs are influenced by their mRNA 3′UTRs. **A.** Workflow for DSG chemical crosslinking in cells, followed by western blot. **B.** Western blot analysis of whole-cell lysates from HeLa cells transfected with HA-ENO1-NU or HA-ENO1-U constructs. Shown are non-crosslinked and DSG crosslinked conditions. 3′UTR-dependent differences in protein crosslinking patterns were quantified using line profiles. The peak corresponding to the expected monomer band is indicated with an arrow. **C.** As in (B) but shown is HA-FOXP2. **D.** Line profiles of the 3′UTR-dependent protein crosslinking patterns of the indicated proteins. **E.** Pearson′s correlation coefficients of the line profiles from the 3′UTR-dependent protein crosslinking patterns from (B-D) are shown. **F.** Model for 3′UTR-dependent folding of IDR-containing proteins. HP clusters in IDRs with folding potential are shown as red dots. Details, see text.

In contrast, when performing crosslinking western blots of protein candidates with HP clusters in their IDRs, including MYC, UTX (encoded by the *KDM6A* mRNA), and FOXP2, in all cases, we observed a 3′UTR-dependent difference in protein crosslinking pattern with Pearson correlation coefficients smaller than 0.7 (Fig. 7C-E, S7H). All these candidates have multivalent HC 3′UTRs (Table S6). This experiment shows that for the tested candidates, the inclusion of 3′UTRs during protein synthesis results in a significant difference in protein crosslinking pattern, which is consistent with a 3′UTR-dependent change in protein state, which most likely occurs during translation (Fig. 7F).

## Discussion

According to Anfinsen’s dogma, the amino acid sequence is sufficient for proper folding of small and fully structured proteins^67^. Our results extend this dogma by showing that the JMJD3 protein, which has hydrophobic amino acid clusters in its IDR only folds properly in the presence of RNA-based IDR chaperone activity, provided by the 3′UTR in the mRNA template. IDR chaperone activity is required for full cellular activity of JMJD3, which acts as histone demethylase. 3′UTRs with IDR chaperone activity often are highly conserved in sequence, are multivalent, and enriched in meshlike cytoplasmic condensates, suggesting that RNA-mediated chaperone activity is spatially restricted in cells.

Based on our validation experiments, where we examined 3′UTR-dependent protein crosslinking patterns of ten candidates, our findings suggest that proteins with hydrophobic amino acid clusters in their IDRs potentially require the presence of RNA acting as IDR chaperone. In contrast, the protein states of fully structured proteins or of proteins that lack hydrophobic clusters in their IDRs were not influenced by including 3′UTRs in the mRNA templates (Fig. 7B-F).

It is currently thought that IDRs do not adopt a single stable conformation in isolation but sample a wide array of distinct conformational states^27^. For 60% of IDR-containing proteins — which lack HP clusters — our data are consistent with this view (Fig. 7F). For proteins with HP clusters in their IDRs, our data suggest that this statement is also true for their post-translational behavior (Fig. 7F). However, during protein synthesis, HP clusters, which only represent 10-15% of sequence in long IDRs, and may bind and potentially interfere with the folding of the structured domains within the same proteins. In the case of JMJD3, we observed that the presence of its corresponding mRNA 3′UTR reduced folding between the IDR and the structured domain and promoted IDR-IDR interactions (Fig. 2H, 2I), consistent with a new function for 3′UTRs acting as co-translational chaperones for IDRs (Fig. 7F).

### Evidence for RNA-based IDR chaperone activity provided by 3′UTRs

Chaperones promote the formation of biologically active conformations and minimize off-pathway reactions. They bind transiently to proteins and are not part of the final folded product. Chaperones bind to unfolded or partially folded proteins, preventing sticky hydrophobic patches from interacting and forming aggregates^68–71^. Whereas traditionally, chaperones were regarded as proteins, more recently it was shown that nucleic acids or polyphosphate also have chaperone activity and reduce protein misfolding and aggregation in vitro^72–76^. Here, we link 3′UTR-dependent cellular activity of transcriptional regulators to 3′UTR-dependent differences in protein conformation, obtained through in-cell crosslinking experiments (Fig. 1J, 1K, 1N, 2I, 2H, 7B-E). Through XL-MS, performed on JMJD3 protein generated in cells from 3′UTR-containing or -lacking mRNA templates, we observed a significantly different crosslinking pattern, providing direct evidence for 3′UTR-dependent protein conformations (Fig. 2H, 2I).

However, the XL-MS experiment does not allow us to distinguish between a direct or indirect effect of the 3′UTR or whether the different protein conformations occur during or after translation. Additional experimental evidence supports a model by which the *KDM6B* 3′UTR directly changes protein folding in a co-translational manner. Through crosslinking experiments, we show that RNA-IDR interaction, provided by in vitro-transcribed 3′UTRs, is necessary and sufficient for a change in protein domain interaction preference in vitro, which fully mimics the domain interactions observed by JMJD3 XL-MS in cells (Fig. 4B, 2H, 2I). Moreover, in cells, both the mRNA 3′UTR as well as the RNA-binding region in the IDR need to be present for a co-translational change in nascent chain interactions (Fig. 4E).

Taken together, these results are consistent with a model by which the 3′UTR binds to the IDR in the JMJD3 nascent chain, thus promoting IDR-IDR interaction, while reducing IDR-folded domain interactions. As a result, the 3′UTR reduces off-pathway conformations that are associated with reduced cellular protein activity. Hydrophobic amino acids at protein domain interfaces have higher sequence conservation^22^, which is consistent with the observed increase in sequence conservation of HP clusters compared with the remaining IDR regions (Fig. 6G). Our data suggest that the mRNA 3′UTR promotes formation of biologically active protein conformations by preventing interdomain folding of hydrophobic amino acid clusters in IDRs during protein synthesis (Fig. 3H, 4F, 7F).

### Difference between post-translational and co-translational RNA binding

Our data suggest that there is a functional difference between post-translational and co-translational RNA-IDR interactions (Fig. 3H). In cells, JMJD3 protein localizes to chromatin, where RNA is abundant. Post-translational RNA-IDR interactions have been widely reported^29,30,33,34,77–79^, which is consistent with our observation that both versions of JMJD3 protein, generated from 3′UTR-containing or -lacking mRNA templates, strongly bind to RNA in cells (Fig. 3E). However, the protein version that was generated from the 3′UTR-containing mRNA template has substantially higher cellular activity than the protein version generated from the NU construct, suggesting that post-translational RNA-binding is not involved in the protein activity regulation investigated here (Fig. 3H). The model that best explains our data suggests that a permanent change in protein folding is caused by co-translational RNA-IDR interaction provided by the 3′UTR. If post-translational RNA-IDR binding could reverse the change in protein folding, we would not observe a significant difference in protein conformation or protein activity, determined by the 3′UTR.

A functional difference for co- and post-translational RNA-protein interaction is further supported by our results on ENO1, where presence or absence or the *ENO1* 3′UTR did not alter protein activity of the fully folded protein (Fig. S1K). However, it was previously shown that post-translational RNA binding to ENO1 protein in stem cells can change its cellular activity^43^. Taken together, our data suggest that co-translational versus post-translational RNA-protein interactions seem to have distinct effects on activity regulation of different protein classes.

### Alternative models by which the *KDM6B* 3′UTR could result in a permanent change in protein folding

Alternative models by which the *KDM6B* 3′UTR could induce a permanent change in protein conformation suggest 3′UTR-dependent recruitment of interaction partners or enzymes that co-translationally bind or modify the JMJD3 nascent chain^6,7^. For JMJD3, this model is unlikely, as we did not observe significant differences in 3′UTR-dependent interactor binding or post-translational modifications, obtained by SILAC-mediated JMJD3 co-immunoprecipitation mass spectrometry analysis. Moreover, the HP region, which is the major negative regulatory domain in the JMJD3 IDR, contains an annotated SUMOylation site. We tested whether modification of this residue could be involved in JMJD3 protein activity regulation. We mutated the residue (K1109R) in the context of the full-length protein and performed IF for H3K27me3 but did not observe a difference in 3′UTR-regulated JMJD3 activity compared to the wildtype (Fig. S3H-K).

Finally, a 3′UTR-dependent co-translational difference in JMJD3 protein conformation could also be caused by a difference in translation rate^8^. Slower translation could promote independent folding of individual domains and prevent folding between domains, whereas fast translation rates could increase the risk of forming kinetically trapped and potentially misfolded intermediates^24^. The testing of five different 3′UTRs in our 3′UTR swapping experiments makes this scenario unlikely, but not impossible. It could be that translation in meshlike condensates decreases translation rates without changing total protein abundance. However, the direct RNA effect on domain interaction preference in vitro and the opposite effects on protein activity seen in the IDR deletion mutants that do not differ in the distribution of optimal and non-optimal codons, makes translation rate-dependent JMJD3 protein folding unlikely (Fig. 3C, 3G, 4B, Table S7)^80,81^.

### Meshlike cytoplasmic condensates seem to emerge as spatial compartments with IDR chaperone activity

With respect to their RNA features, we observed that HC 3′UTRs are multivalent, a physical RNA feature that is necessary for entrapment in meshlike cytoplasmic condensates (Fig. 5F, 5H)^12^. In addition to multivalent 3′UTRs, we found that the presence of condensate-enriched RNA-binding proteins, such as TIAL1, is also required for 3′UTRs to act as IDR chaperones (Fig. 5K)^13^. Moreover, IDR chaperone activity correlates with mRNA localization to meshlike condensates, as mRNAs that poorly localize to meshlike condensates do not have JMJD3 IDR chaperone activity (Fig. 5B, 5H). Taken together, these data are consistent with a model by which meshlike condensates are cytoplasmic compartments that provide favorable folding environments for IDR-containing proteins, as they are enriched for mRNAs whose 3′UTRs have IDR chaperone activity.

However, direct evidence for 3′UTR-dependent protein folding happening during translation in meshlike condensates is currently lacking. Moreover, the molecular characteristics of RNAs capable of acting as IDR chaperones are unknown, as is the role of RNA-binding proteins in this process. It is not known if RNA motifs, responsible for mRNA localization to meshlike condensates and motifs that mediate IDR chaperone activity are overlapping or distinct. Although mRNAs with HC 3′UTRs are strongly enriched in meshlike condensates (Fig. 5E), our study did not reveal why these 3′UTRs have hundreds of conserved nucleotides. Lastly, it is unclear if IDR chaperone activity is performed by the 3′UTR in the mRNA that encodes the IDR-containing protein or whether condensate-enriched 3′UTRs can act in trans on IDRs of other nascent chains.

As the 3′UTRs of condensate-enriched mRNAs have weak predicted structures and have features that can be interpreted as structural plasticity (Fig. S5J), we speculate that these RNA features could be necessary for RNAs acting as IDR chaperones, thus making them conditionally disordered, a characteristic that is typical for protein chaperones^82^. However, biophysical experiments are needed for a better characterization of the RNA features responsible for IDR chaperone activity.

### Implications for ectopic gene expression studies

The importance of 3′UTRs in mRNA templates that encode proteins with long IDRs is suggested by their extensive sequence conservation (Fig. 1A-D). HC 3′UTRs often are as conserved as their corresponding CDS, suggesting a critical role in protein biogenesis. Proteins with long IDRs and hydrophobic clusters are substantially enriched in transcription factors and chromatin regulators (Fig. 1E)^35,40,83^.

Our work has widespread and important implications for the study of IDR-containing proteins using vector constructs. Ectopic gene expression is a widely used tool to study protein functions in cells, transgenic animals, and through recombinant proteins. In nearly all cases, viral or cDNA expression vectors are used that drive expression of the CDS only. This has mostly historical reasons as vector technology was developed in the 1970s, when it was unknown that approximately 50% of mRNA sequence is contributed by 3′UTRs^1,2,84^. Using CDS-only vectors for expression of IDR-containing proteins may generate only partially active proteins due to the lack of IDR chaperones. Moreover, a recombinantly generated protein is already in a ‘post-translational state’. The lack of IDR chaperones during protein synthesis may kinetically trap these proteins in non-functional conformations, thus potentially generating misleading results. As IDRs are widespread in the human proteome^26,27,35,83^, new experimental approaches need to be established that include 3′UTRs in mRNAs and IDRs in proteins when studying protein functions to fully incorporate their regulatory potential.

### Limitations of the study

Although in vitro, RNA-IDR interaction is necessary and sufficient for a change in JMJD3 domain interaction preference, which mimics the protein folding change of cellular, full-length JMJD3 observed by XL-MS, we did not directly show that the *KDM6B* 3′UTR binds to the JMJD3 nascent chain while being translated in meshlike condensates.

Although, we present several lines of evidence that 3′UTRs can act as IDR chaperones, we have not directly ruled out that the *KDM6B* 3′UTR could also influence translation rate with subsequent consequences on protein folding.

Chaperones are molecules that transiently interact with their substrate proteins but are not part of the final product. Although, we think it is unlikely that 3′UTRs are part of the final protein products, this part of the model has not been tested directly.

## Acknowledgements

This work was funded by a grant from the Pershing Square Foundation, William Ackman, and Neri Oxman, a grant from the G. Harold and Leila Y. Mathers Foundation, the NIH Director’s Pioneer Award (DP1GM123454) and the NIH grant R35GM144046 to C.M. as well as the MSK Core Grant (P30 CA008748). We thank Ting Cai for providing the *KDM6B* 3′UTR deletion iPS cells and Ellen Horste for generating the HeLa *TIAL1* knockout cells and help with RNA-FISH. We thank Mervin Fansler for early work on 3′UTR conservation and help with RNA-seq analysis, Chiara Evans for help with the co-translational disome assay, and Jiaxiang Ren for providing code for the analysis of immunofluorescence images. We thank Mara Monetti and Zhuoning Li from the Proteomics Core at MSKCC for performing and analyzing the mass spectrometry experiments and for useful discussions. We thank Pierre-Jacques Hamard and Richard Koche from the Epigenetics Research Innovation Lab at MSKCC for performing and analyzing the ChIP-seq experiments. The ChIP-seq samples were sequenced at the MSKCC Integrated Genomics Operation Core facility. We thank Ting Cai and all members of the Mayr lab, as well as Thomas R. Cech (University of Colorado Boulder), Nancy M. Bonini (University of Pennsylvania), and Emmanuel Levy (University of Geneva) for critical reading of the manuscript and helpful discussions.

## Author contributions

Y.L. performed all experiments using vector constructs in cells. The in vitro RNA binding and crosslinking assays were performed and analyzed by Y.Z. M.C.W performed the experiments on endogenous *KDM6B* 3′UTR deletion. S.B. performed the computational analyses with input from C.M. Y.L. and C.M. conceived the project, designed the experiments, and wrote the manuscript with input from all authors.

## Declaration of interest

C.M. is a member of the *Cell* advisory board.

The authors have no other competing interests to declare.

## Supplementary Figure legends

**Figure S1.**
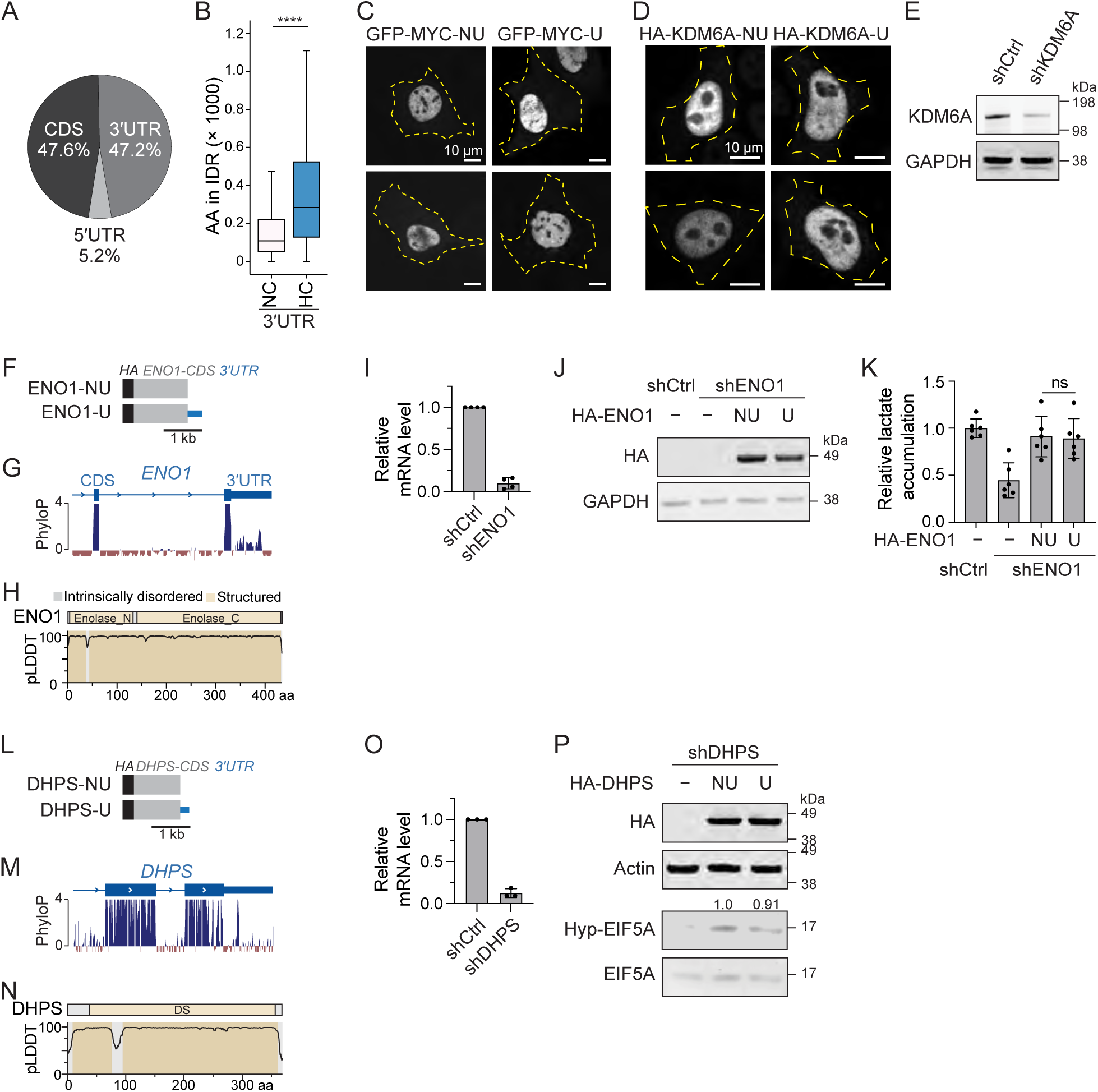
NC 3′UTRs in mRNA templates do not alter the activity of proteins encoded by the mRNA, related to Figure 1. **A.** Proportion of nucleotides assigned to 5′UTRs, CDS, and 3′UTRs in human mRNAs (*N* = 19,556)^2^. **B.** As in Fig. 1C but shown is the number of residues in IDRs based on classification using pLDDT scores from AlphaFold2. IDRs are defined as pLDDT < 80. Mann-Whitney test, *P* = 5.5 x 10^-248^. **C.** Live-cell fluorescence microscopy of GFP-MYC-NU or GFP-MYC-U in U2OS cells. Representative image is shown. The yellow dotted line indicates the cell outline. MYC protein localizes to the nucleus in a 3′UTR-independent manner. **D.** As in (C), but anti-HA immunofluorescence staining of transfected HA-KDM6A-NU or HA-KDM6A-U in MCF7 cells. **E.** Western blot of endogenous UTX/KDM6A levels in MCF7 cells stably expressing either a control shRNA or an shRNA targeting KDM6A. GAPDH was used as loading control. **F.** cDNA vector constructs used for ENO1 activity assays in cells, showing the HA-tag (black), CDS (gray) and full-length 3′UTR (blue). **G.** PhyloP100way track of the last two exons of the *ENO1* gene. **H.** Prediction of ENO1 structured domains (pLDDT ≥ 80, brown) and IDRs (pLDDT < 80, gray) using AlphaFold2 pLDDT scores. Names of annotated protein domains are given. **I.** Quantification of shRNA-mediated ENO1 knockdown by qRT-PCR. Data are normalized to *RPLP0* and are represented relative to cells transduced with a scrambled shRNA (shCtrl). Data and error bars indicate mean ±Lstd of 4 biological replicates. **J.** Western blot analysis after transfection of constructs shown in (F) into HeLa cells depleted of ENO1. GAPDH was used as a loading control. **K.** Comparison of the cellular activity of ENO1 expressed from NU or U constructs in HeLa cells, indirectly measured by the accumulation of lactate in the medium. Data and error bars indicate mean ±Lstd of 6 biological replicates. Two-tailed t-test: ns. **L.** As in (F), but DHPS expression constructs are shown. **M.** As in (G), but the PhyloP track of DHPS is shown. **N.** As in (H), but DHPS pLDDT values and domains are shown. **O.** As in (I), but the knockdown efficiency of DHPS in HeLa cells was analyzed. **P.** Comparison of DHPS cellular activity by measuring EIF5A hypusination levels via western blot. Endogenous DHPS was knocked down in HeLa cells using shRNA to evaluate the effect of transfected DHPS constructs. The relative EIF5A hypusination level was calculated by normalizing the hypusination band intensity to total EIF5A protein and is displayed above the blot.

**Figure S2.**
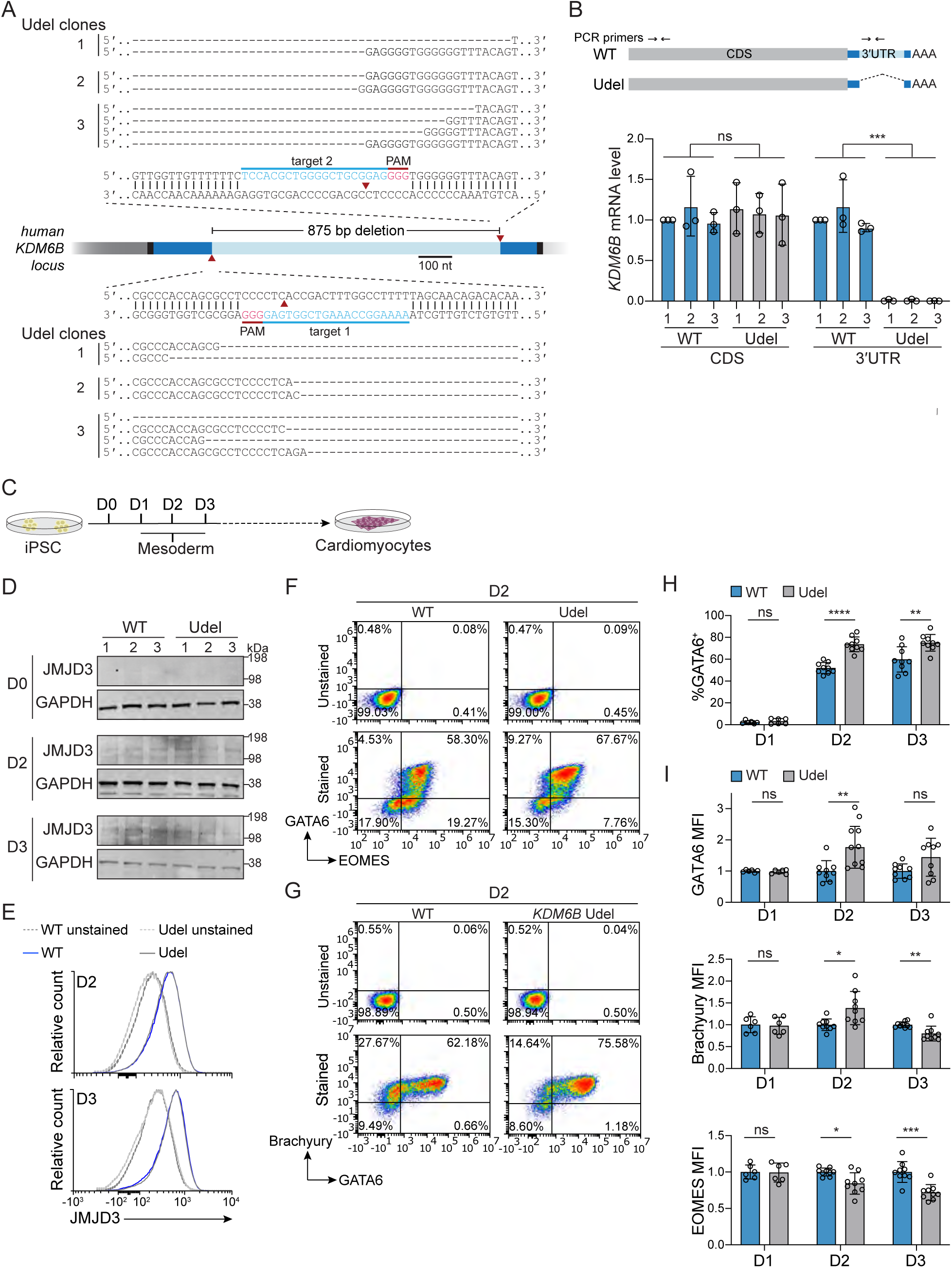
Partial deletion of the endogenous *KDM6B* 3′UTR affects mesendoderm differentiation, related to Figure 1. **A.** Sequences of the human *KDM6B* locus targeted by Cas9 and guide RNAs. The genomic sequence deleted in *KDM6B* 3′UTR deletion cells (Udel) is shown in skyblue. sgRNA target sites and PAMs are indicated by blue and magenta bars, respectively, and predicted cutting sites are marked by a red triangle. Sequence alignment of KDM6B alleles spanning the deletion sites in Udel iPSC cell clones obtained from amplicon sequencing are shown. **B.** *KDM6B* mRNA expression measured by qRT-PCR with a primer pair located in the CDS or 3′UTR in the indicated samples derived from iPSC cells. Data are shown as mean ±Lstd of 3 biological replicates. WT, wild type. T-test for independent samples: ***, *P* < 0.001; ns, not significant. **C.** Schematic for in vitro cardiomyocyte differentiation using the GiWi protocol. The cells adopt mesoderm and mesendoderm markers at d2 and d3, which was analyzed here. **D.** Western blot of endogenous JMJD3 protein in WT and Udel iPSC clones at day 0, day 2, and day 3 of the differentiation protocol. GAPDH was used as the loading control. **E.** Representative flow cytometry analysis of endogenous JMJD3 protein expression WT and Udel iPSC clones at day 2 and day 3 of the differentiation protocol. **F.** Representative flow cytometry plots at day 2 of the differentiation protocol, analyzing the percent of GATA6- and EOMES-positive cells in the indicated samples. **G.** As in (F), but the percent of Brachyury- and GATA6-positive cells is shown. **H.** Quantification of day 1-3 flow cytometry data for percent GATA6-positive cells is shown as mean ± std of *N* = 3 biological replicates for each of the three WT and Udel clones. Two-tailed t-test, ns, not significant; **, *P =* 0.005; ****, *P* < 0.00001. **I.** As in (H), but mean fluorescence intensity (MFI) as measure of protein expression of GATA6, Brachyury, and EOMES is shown as mean ± std obtained from 3 biological replicates each for three WT and three Udel clones. Two-tailed t-test: ns, not significant; *, *P* < 0.05; **, *P* < 0.01; ***, *P* < 0.001.

**Figure S3.**
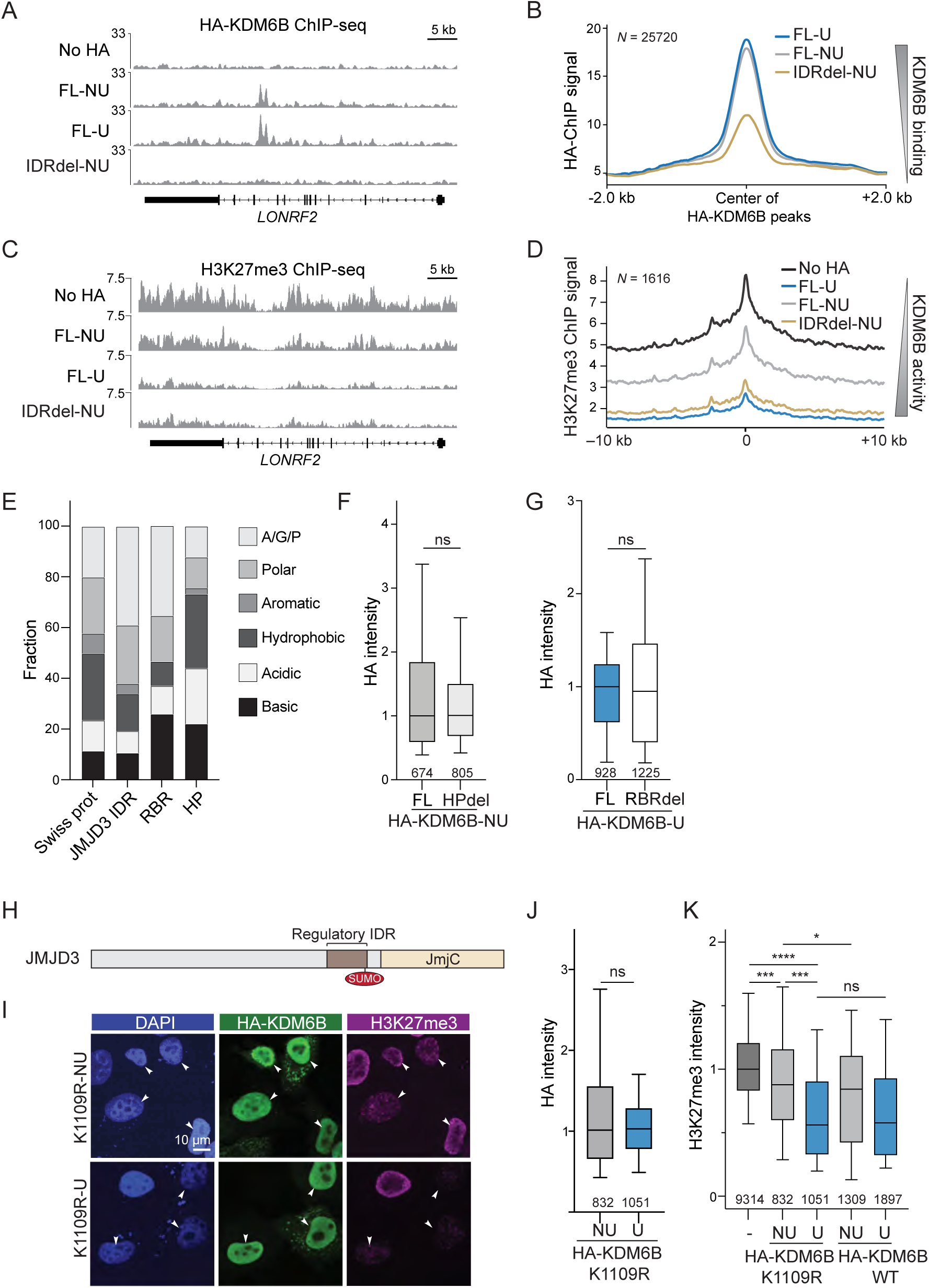
The JMJD3 IDR mediates 3’UTRIZIdependent activity regulation independently of its roles in chromatin occupancy and SUMOylation, related to Figure 2 and 3. **A.** Genome browser view of genomic context of *LONRF2* for anti-HA ChIP-seq signal (depth-normalized read count of one of the replicates) is shown. ChIP-seq was performed after transfection of the indicated KDM6B constructs, shown in Fig. 1H and 2B. No HA indicates negative control. **B.** Metaplot analysis of experiment from (A) using all anti-HA ChIP-seq peaks with *P* < 0.001. **C.** As in (A), but spike-in normalized anti-H3K27me3 ChIP-seq signal (depth-normalized read count of one of the replicates) is shown. **D.** Metaplot analysis of experiment from (C) using all anti-H3K27me3 ChIP-seq peaks with *P* < 0.001 that showed at least a 50% signal reduction in KDM6B-U transfected samples compared to the control. **E.** Stacked bar chart showing the relative fractions of each amino acid category across all proteins in the Swiss-Prot database, the JMJD3 IDR, the RBR, and the HP region, respectively. **F.** As in Fig. 1M, but JMJD3 protein abundance of full-length and HPdel was measured as HA intensity from *N* = 3 independent experiments. Mann-Whitney test: ns, not significant. **G.** As in Fig. 1M, but JMJD3 protein abundance of full-length and RBRdel was measured as HA intensity from *N* = 3 independent experiments. Mann-Whitney test: ns, not significant. **H.** Schematic representation of the annotated SUMOylation site within the regulatory IDR. To examine if 3′UTR regulates JMJD3 activity via SUMOylation at this site, K1109 was mutated into R to abolish SUMOylation while maintaining the positive charge. **I.** As in Fig. 1L, but JMJD3 cellular activity in HeLa cells after transfection of the K1109R mutant constructs. **J.** As in Fig. 1M, but JMJD3 protein abundance of the K1109R constructs was measured as HA intensity from *N* = 3 independent experiments. Mann-Whitney test: ns, not significant. **K.** As in Fig. 1N, but the cellular activity of wildtype JMDJ3 and K1109R were measured as reduction in H3K27me3. Normalized H3K27me3 intensity values obtained from *N* = 3 independent experiments. Mann-Whitney test: ****, *P* < 10^-204^; ***, *P* < 10^-24^; *, *P* < 10^-6^; ns, not significant.

**Figure S4.**
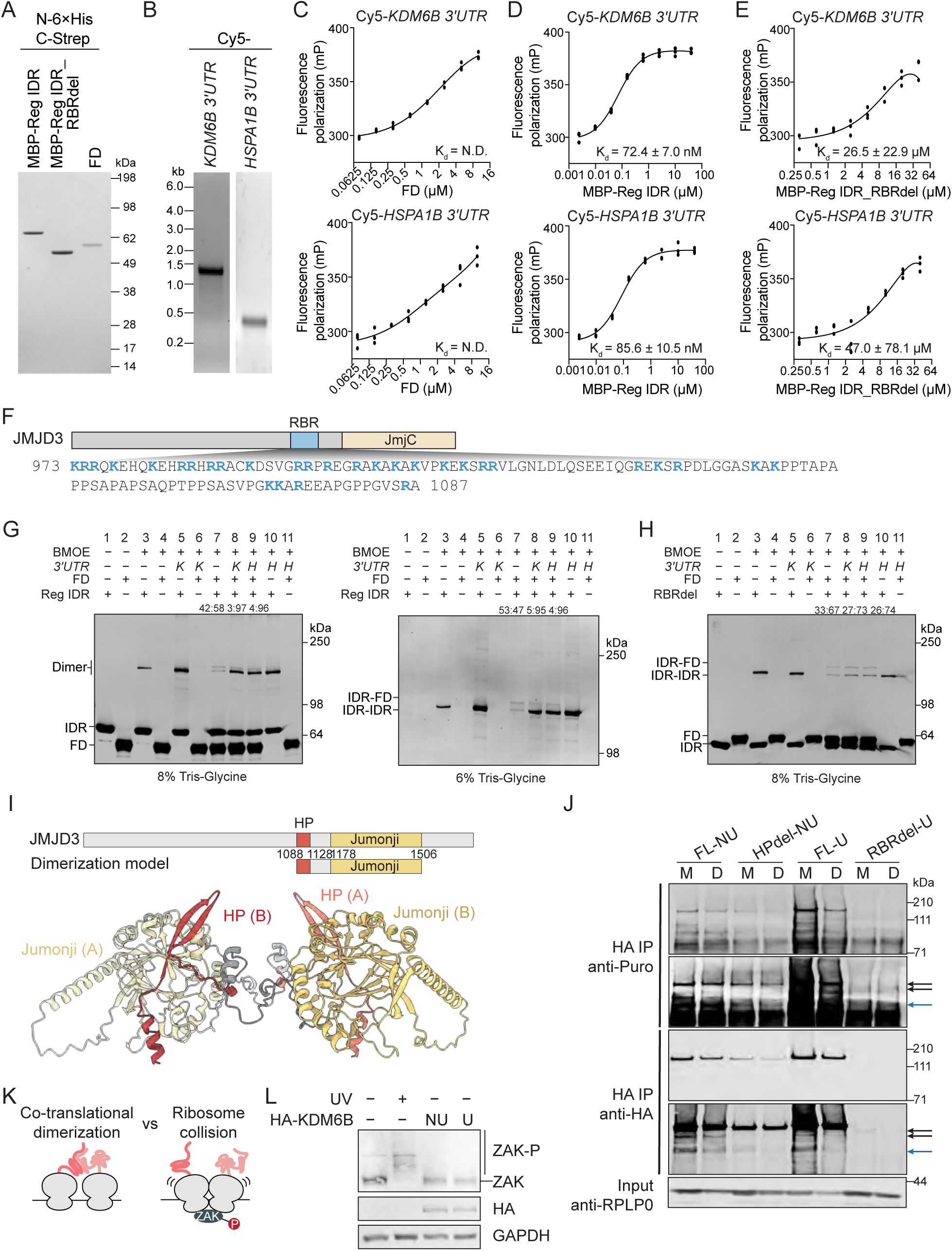
RNAs bind the JMJD3 IDR, regulating inter-domain interactions in vitro and controlling co-translational JMJD3 dimerization in cells, related to Figure 4. **A.** Coomassie-stained SDS–PAGE showing recombinant MBP-tagged regulatory IDR, MBP-tagged RBR-deleted regulatory IDR (RBRdel), and folded domain (FD) from JMJD3. All proteins contain a N-terminal 6xHis and a C-terminal Strep tag II. **B.** Agarose gel showing in vitro transcribed Cy5-labeled *KDM6B* (1164 nt) and *HSPA1B* 3′UTRs (378 nt). The first 110 nt of the *KDM6B* 3′UTR are GC-rich and were removed to allow transcription. **C-E.** Fluorescence polarization assay to measure RNA-protein interactions. Top: As RNA, the Cy5-labeled *KDM6B* 3′UTR (1164 nt) was used. Bottom: As RNA, the *HSPA1B* 3′UTR (378 nt) was used. Fluorescence polarization values obtained from *N* = 3 independent experiments are shown. The values were fitted to a sigmoidal binding curve, and the K_d_ was calculated in Prism using the one-site binding model. The dissociation constant (K_d_) is reported as ± error of fitting. **F.** Amino acid sequence of the RNA-binding region (RBR) in JMJD3. Basic residues are highlighted in blue. **G.** The uncropped blot showing RNA-dependent protein-protein interactions generated through thiol-thiol crosslinking between recombinant JMJD3 regulatory IDR and FD by BMOE-mediated thiol-thiol crosslinking. The same reactions were resolved on either an 8% (left) or 6% (right) Tris-Glycine gel to allow clear visualization of individual monomer and dimer bands, respectively. *K*, *KDM6B* 3′UTR was used; *H*, *HSPA1B* 3′UTR was used as RNA. **H.** As in (G) but showing the crosslinking experiment between RBRdel and FD. **I.** AlphaFold3 prediction of full-length JMJD3 homodimer but shown are only the interacting domains (aa 1086-1510), color-coded as in the protein model shown above. Red, HP region; brown, Jumonji domain. (A) and (B) indicate two different JMJD3 molecules. **J.** Representative immunoblots of the co-translational disome assay. Shown is the abundance of newly synthesized nascent chains derived from various HA-tagged KDM6B constructs in either the monosome (M) or disome (D) fractions obtained by sucrose gradient centrifugation after their transfection into HeLa cells. As input, RPLP0 in each M or D fraction was blotted and used for normalization purposes. Puro, puromycin. Black arrows indicate the nascent chains that overlap with the full-length HA-KDM6B bands, which were used for quantification. The blue arrow indicates a lower molecular weight, HA-tagged nascent chain that displayed a similar trend as the full-length band. **K.** Disomes can form from interactions between nascent chains (left) or from ribosome collisions induced by stress (right). **L.** Western blot of Phos-tag gels for ZAK phosphorylation in HeLa cells subjected to the indicated treatment.

**Figure S5.**
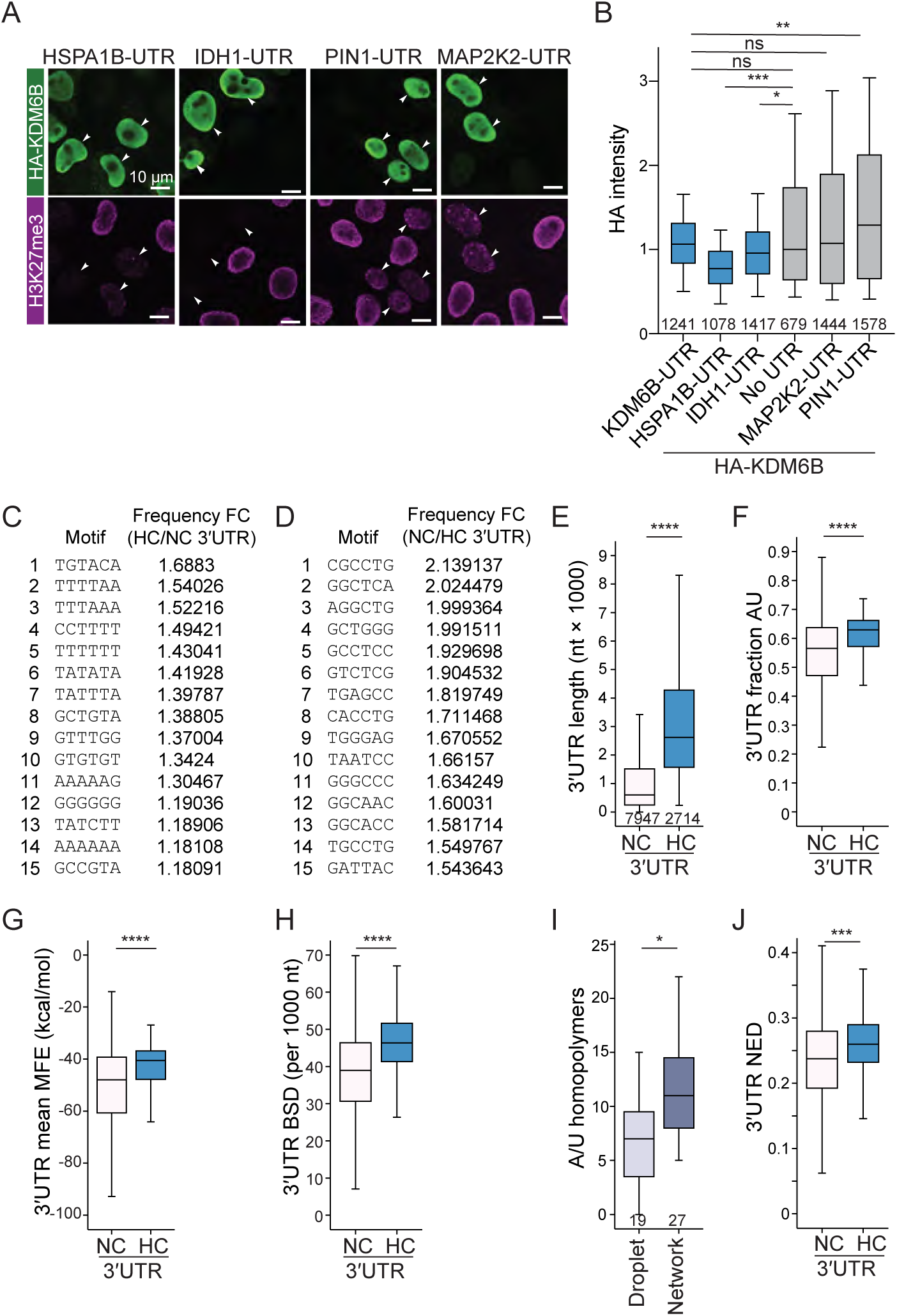
HC 3′UTRs are multivalent, a feature that correlates with IDR chaperone activity, related to Figure 5. **A.** Immunofluorescence assay to assess 3′UTR-dependent JMJD3 activity in HeLa cells after transfecting the indicated 3′UTR-swapped constructs. **B.** Quantification of transfected JMJD3 abundance as HA intensity values from (A). Shown are boxplots generated from *N* = 3 independent experiments. Mann-Whitney test: ns, not significant; *, *P =* 1.3 x10^-5^; **, *P* = 6.9 x 10^-15^; ***, *P =* 1.2 x 10^-29^. **C.** The top 15 6-mer motifs significantly enriched in HC vs. NC 3′UTRs. **D.** As in (C), but motifs enriched in the NC compared to HC 3′UTRs are shown. **E.** 3′UTR length distribution of human mRNAs in HC 3′UTRs (*N* = 2714) vs NC 3′UTRs (*N* = 7947), Mann-Whitney test, *P* = 0. **F.** As in (E), but AU content in 3′UTRs is shown. Mann-Whitney test, *P* = 2.6 x 10^-130^. **G.** As in (E), but mean minimum free energy (MFE) of 3′UTRs is shown. Mann-Whitney test, *P* = 8.5 x 10^-106^. **H.** As in (E), but RNA-RNA binding site density in 3′UTRs is shown. Mann-Whitney test, *P* = 3.6 x 10^-228^. **I.** A/U homopolymers in 3′UTRs that induced droplet-like (*N* = 19) or meshlike condensates (*N* = 27) in vitro, obtained from Ma et al., Elife (2021)^12^. Mann-Whitney test, *P* = 0.018. **J.** As in (E), but normalized ensemble diversity (NED) values of the 3′UTRs are shown. Mann-Whitney test, *P* = 4.9 x 10^-68^.

**Figure S6.**
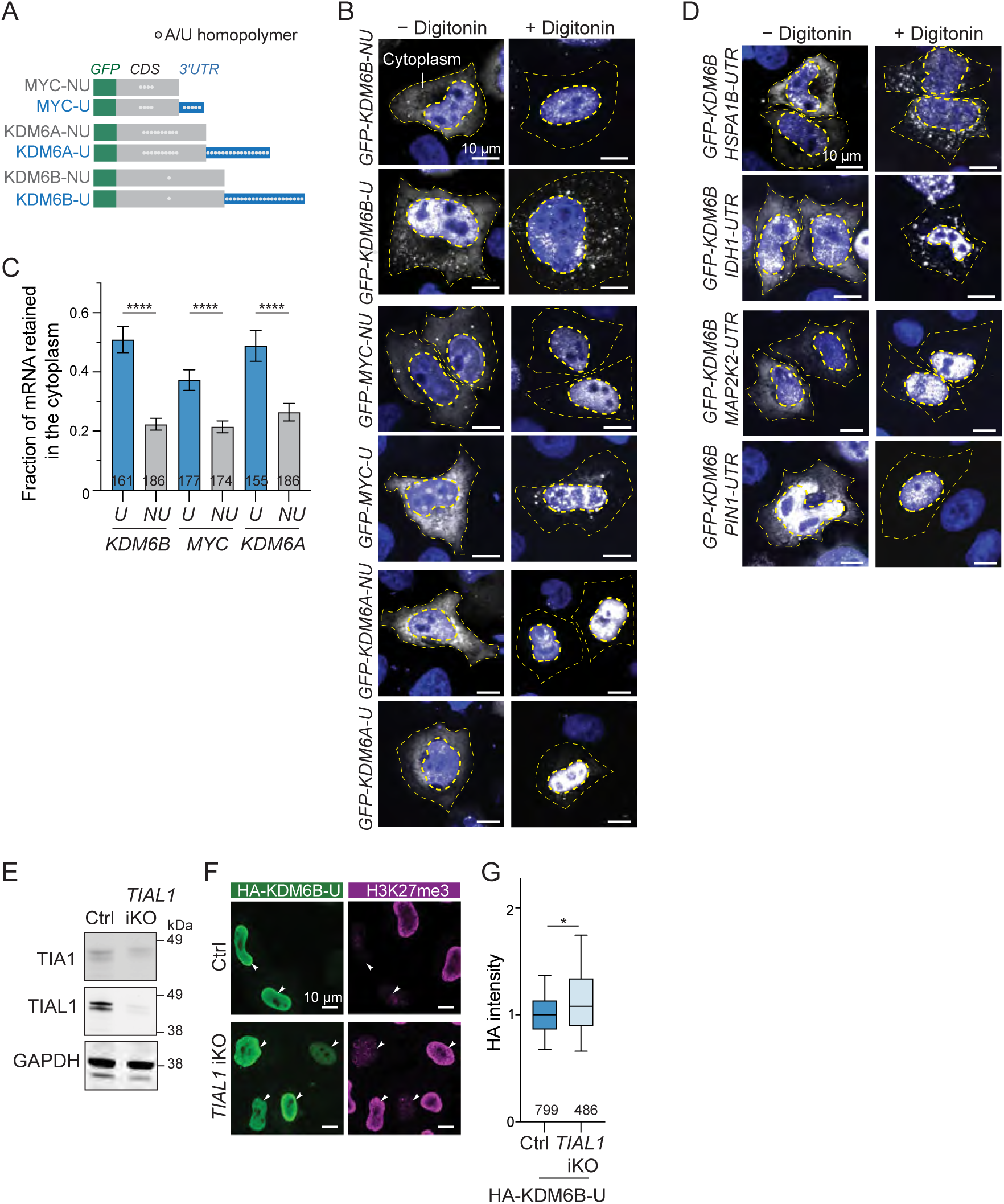
Multivalent 3′UTRs are necessary for mRNA enrichment in meshlike condensates, related to Figure 5. **A.** GFP-tagged constructs used for RNA-FISH. The white dots indicate A/U homopolymers and serve as surrogate marker for RNA valency. **B.** RNA-FISH (white) of mRNAs from the indicated transfected constructs in HeLa cells before (-) and after (+) digitonin extraction. DAPI (blue) stains nuclei. Thin yellow dashed lines indicate cell boundaries, and thick dotted lines mark the nucleus. As digitonin only extracts cytoplasmic mRNAs, only cytoplasmic RNA-FISH signal was analyzed. Representative images are shown. **C.** Quantification of (B). Fraction of digitonin-resistant mRNA in the cytoplasm shown as mean ± SEM from *N* = 3 independent experiments. The number of analyzed cells is shown. Mann-Whitney test: ****, *P* < 0.0001. **D.** As in (B), but RNA-FISH was performed on *KDM6B* mRNA with swapped 3′UTRs. Constructs shown in Fig. 5A. **E.** Western blot showing TIAL1 expression in a doxycycline inducible CRISPR/Cas9 HeLa cell line expressing either gRNAs targeting TIAL1 (iKO) or non-targeting control (Ctrl). GAPDH was used as the loading control. **F.** As in Fig 1L, but JMJD3 cellular activity in Ctrl or *TIAL1* iKO HeLa cells after transfection of the HA-KDM6B-U construct. **G.** As in Fig. 1M, but JMJD3 protein abundance of transfected full-length JMJD3 in the indicated cell lines from *N* = 3 independent experiments is shown. Mann-Whitney test: *P* = 1.9 × 10^-10^.

**Figure S7.**
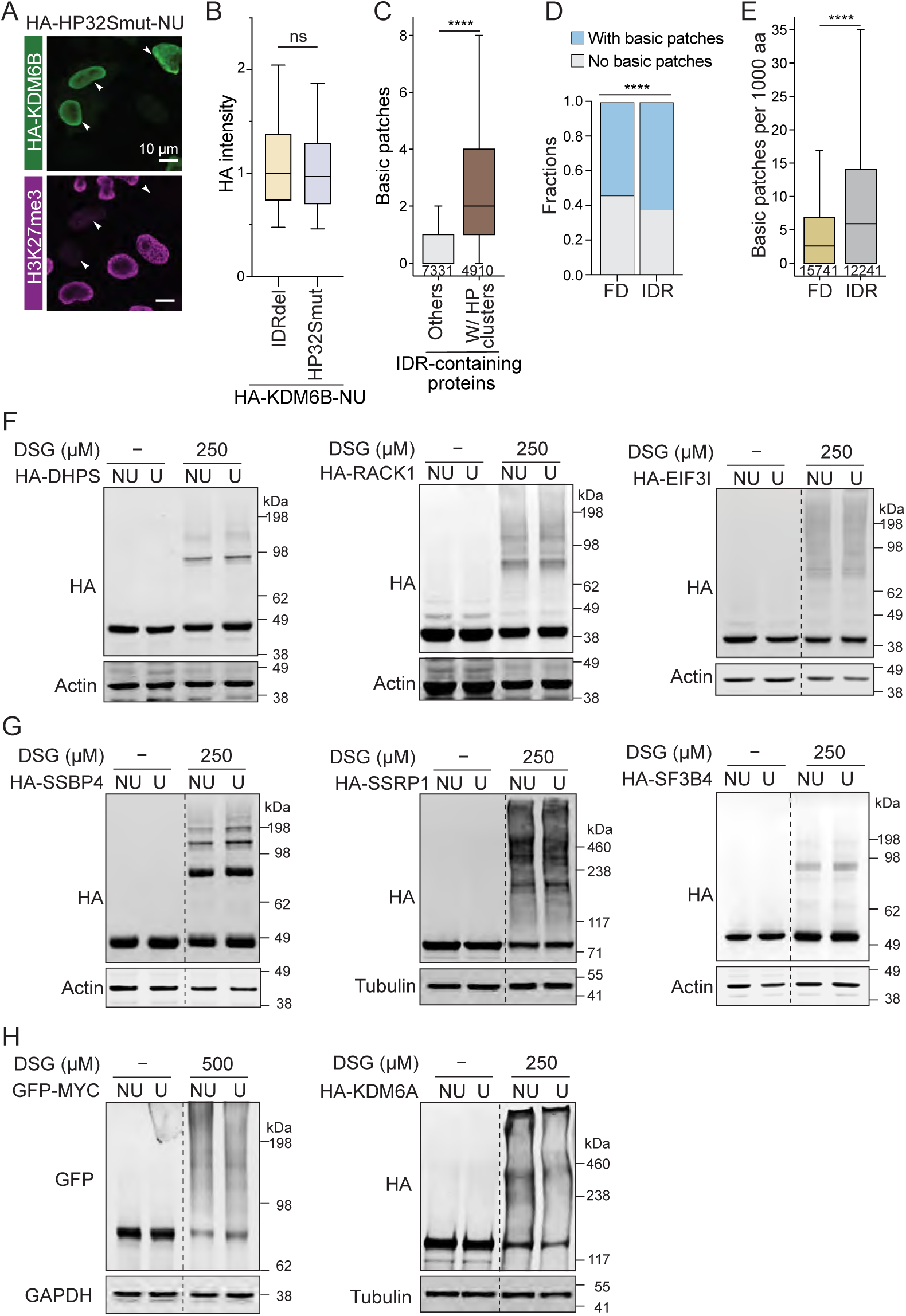
Protein crosslinking patterns of IDR-containing proteins with hydrophobic clusters change in a 3′UTR-dependent manner, related to Figure 6 and 7. **A.** As in Fig. 1L, but JMJD3 cellular activity in HeLa cells after transfection of the HP32Smut construct. **B.** As in Fig. 1M, but JMJD3 protein abundance of the IDRdel and HP32Smut constructs was measured as HA intensity from *N* = 3 independent experiments. Mann-Whitney test: ns, not significant. **C.** As in Fig. 6E, but the number of basic patches is shown. IDR-containing proteins *N* = 12241, IDRs with HP clusters, *N* = 4910, and IDRs without HP clusters, *N* = 7331. Mann-Whitney test: ****, *P* = 0. **D.** Fraction of folded domains with basic patches (8465/15741) and fraction of IDRs with basic patches (7580/12241). Test for independent proportions, ****, *P* < 0.0001. **E.** Basic patches per 1000 amino acids in the longest folded domains (FD, *N* =15741) vs the longest IDRs (*N* = 12241) in each protein. Mann-Whitney test: *P* = 1.6 x 10^-236^. **F-H.** Protein crosslinking patterns in cells after transfection of the indicated HA-tagged constructs were analyzed by DSG cross-linking and western blot analysis. **F.** DSG crosslinking western blot of HA-tagged folded proteins shows 3′UTR-independent protein crosslinking patterns. Actin was used as loading control. **G.** As in (F) but shown are HA-tagged IDR-containing proteins free of HP clusters. Actin or Tubulin were used as loading controls. **H.** DSG crosslinking western blot of HA-tagged proteins containing IDRs with HP clusters shows 3′UTR-dependent protein crosslinking pattern. GAPDH or Tubulin were used as loading controls.

## Supplementary Tables

**Table S1. mRNA and protein features of human protein coding genes, related to Figure 1, 6, and 7.**

**Table S2. RNA-seq analysis of 3′UTR-regulated MYC activity, related to Figure 1J**.

**Table S3. RNA-seq analysis of 3′UTR-regulated UTX transcriptional targets, related to Figure 1K**.

**Table S4. XL-MS data of cellular JMJD3 translated from NU or U constructs, related to Figure 2**.

**Table S5. Spearman’s correlation coefficients across different RNA features, related to Figure 5**.

**Table S6. mRNA and protein features and expression levels of the ten candidates experimentally tested for 3′UTR-dependent protein states, related to Figure 7**.

**Table S7. Codon content of the wild-type and mutant JMJD3 mRNAs, related to Figures 1,3, and 6.**

**Table S8. Sequences of oligonucleotides used, related to Key Resources Table.**

**Table S9. Statistical test values, related to Figures 1-7**.

## STAR methods

### RESOURCE AVAILABILITY

#### Lead contact

Further information and requests for resources and reagents should be directed to and will be fulfilled by the Lead contact, Christine Mayr.

#### Materials availability

- All plasmids generated in this study will be deposited to Addgene.
- The gene edited cell lines HeLa TIAL1 iKO, iPSC 731.2B KDM6B Udel clones and corresponding ctrl clones generated in this study are available from the Lead Contact with a completed Materials Transfer Agreement.

#### Data and code availability

Calculated RNA and protein parameters are reported in Table S1. The processed RNA-seq data for MYC and UTX/KDM6A are reported in Tables S2 and S3 and their raw data are accessible through GEO accession number GSE301135. The JMJD3/KDM6B HA and H3K27me3 ChIP seq data are accessible through GEO accession number GSE301136. The data of the XL-MS experiment have been deposited in the PRIDE repository with the dataset identifier PXD072448. The processed data are reported in Table S4. The correlation analysis to identify a surrogate marker for RNA valency is reported in Table S5. The protein and RNA features of the experimentally validated proteins are reported in Table S6. The codon usage ratio of our KDM6B constructs is reported in Table S7.

### Experimental model

#### Cell lines

HeLa (human cervical cancer line, female origin) was previously described^11^. MCF7 (human breast cancer, female origin) and U2OS (human osteosarcoma line, female origin) were previously described^5^. All cells were maintained at 37°C with 5% CO2 injection in Dulbecco’s Modified Eagle Medium (DMEM) containing 4.5 g/l glucose, 10% heat inactivated fetal bovine serum, 100 U/ml penicillin and 100 μg/ml streptomycin.

##### Doxycycline-inducible *TIAL1* knockout HeLa cells (*TIAL1* iKO)

Doxycycline (Dox)-inducible Cas9 (iCas9) HeLa Cells were generated as previously described^13^. The iCas9 line was transduced with a lentiviral construct harboring a pair of guide RNAs either targeting *TIAL1* or non-targeting control sgRNAs. The plentiGuide-puro vector was adapted to generate these constructs, where synthetic DNA fragments were ordered from Genewiz/Azenta and inserted. The guide RNA sequences are listed in Table S8. Lentivirus was generated in HEK293T cells, and iCas9 cells were transduced followed by puromycin selection. For induction of gene knockouts, *TIAL1* iKO and control cells were treated with 100 ng/ml Dox for five days, after which TIAL1 and TIA1 expression were evaluated by western blotting.

##### iPSCs with partial *KDM6B* 3′UTR deletion

The 731.2B iPSCs were obtained from the Stem Cell Core facility at MSKCC and cultured using a StemFlex medium (Thermo Fisher Scientific, A3349401) on a Matrigel (Corning, 354277)-coated surface in a humidified 37°C incubator with 5% CO2. For general purposes, the cells were detached with 0.5 mM EDTA. To conduct gene editing through CRISPR-Cas9 system, the sgRNAs were designed using CRISPOR and ordered from Synthego with the modification of 2’-O-methyl analogs and 3’ phosphorothioate linkages. The control sgRNAs were designed without human genome binding capacity, whereas two sgRNAs were used to generate a genomic deletion that corresponds to the *KDM6B* 3′UTR at the mRNA level. The iPSCs were detached with TrypLE Select and replated to a 12-well at 250 000 cells per well before performing sgRNA transfection. Lipofectamine RNAiMax (Thermo Fisher Scientific, 13778150) was used to deliver sgRNAs one day after the induction of Cas9 under 1 µg/ ml Doxycycline (Acros Organics, 446060050) in Stemflex media. After two days of gene editing, the cells were detached with Accutase and passed through a 40 µm strainer to make a single cell suspension for colony formation. To increase cell viability, the cells were plated under 10 µM Y27632 in Stemflex media and washed out on the second day. Gene editing of single colonies was validated by Amplicon Sequencing (Genewiz) and qRT-PCR analysis for *KDM6B* CDS and 3′UTR mRNA expression in three 3′UTR deletion (Udel) and three control clonal cell lines. The primers are listed in Table S8.

## Method details

### Vector constructs

#### GFP-tagged constructs

For GFP-tagged constructs, pcDNA3.1-puro mGFP (monomeric GFP, A207K), which was previously reported^11^, was used. To generate GFP-MYC, the *MYC* promoter (2780 bp; hg38, chr8:127733806-127736585) was PCR-amplified from HeLa genomic DNA and inserted between *BglII* and *NheI* sites. To generate GFP-MYC-NU, the *MYC* coding sequence (1365 bp) was PCR-amplified from HeLa cDNA and inserted between *BsrGI* and *BamHI* sites. For GFP-MYC-U, the *MYC 3*′*UTR* (473 bp) was amplified and inserted between *BamHI* and *EcoRI* of the NU construct.

To obtain GFP-KDM6A and GFP-KDM6B constructs for RNA-FISH, the coding sequences were PCR-amplified from HA-tagged constructs (see below) and inserted into the pcDNA3.1-puro mGFP vector between *BsrGI* and *NotI* or *KpnI* sites, respectively. The 3′UTRs were then inserted between *NotI* or *KpnI* and *XhoI* sites.

#### HA-tagged constructs

KDM6A (NM_001291416) and KDM6B (NM_001080424) expression constructs in the pCMV-HA backbone were obtained from Addgene (#24168 and #24167)^45^ and used as NU constructs. For the U constructs, the full-length *KDM6A 3*′*UTR* (1368 bp) and full-length *KDM6B 3*′*UTR* (1274 bp) were each amplified from HeLa genomic DNA and inserted between *NotI* and *XhoI* restriction sites. For ChIP-seq, a plasmid containing NLS-KDM6Bs (amino acids 1027–1682)^45^ was used as IDRdel. The NLS (PKKKRKV) was derived from SV40 large tumor antigen and was added to the N-terminus via PCR. For all other assays, IDRdel contains this NLS at the N-terminus and amino acids 1141-1682.

All other HA-tagged expression constructs were cloned into the pCMV-HA backbone using similar approaches and were used in the chemical crosslinking experiments. For NU constructs, coding sequences (CDS, lacking a start codon) of ENO1 (1302 bp, NM_001428.5), SSRP1 (2127 bp, NM_003146), SF3B4 (1272 bp, NM_005850), SSBP4 (1155 bp, NM_032637), EIF3I (975 bp, NM_003757), DHPS (1107 bp, NM_001930) and RACK1 (951 bp, NM_006098) were PCR amplified from HeLa cDNA and inserted between *KpnI* and *NotI,* except for the FOXP2 coding sequence (2145 bp, NM_014491), which was PCR-amplified from HEK293 cDNA. The FOXP2-NU construct also includes a C-terminal FLAG-tag (DYKDDDK), added via PCR. For the FOXP2-U construct, the full-length *FOXP2 3*′*UTR* (3838 bp) was inserted between *NotI* and *XhoI* sites. For the U constructs of low-valency mRNAs, the coding sequences together with the 3′UTRs (ENO1, 1662 bp; SSRP1, 2546 bp; SF3B4, 1497 bp; SSBP4, 1471 bp; EIF3I, 1383 bp; DHPS, 1228 bp; RACK1, 1030 bp) were directly amplified from cDNA and inserted between *KpnI* and *NotI*.

For chimeric constructs containing the KDM6B CDS and various 3′UTRs, 3′UTR sequences (*IDH1 3*′*UTR*, 850 bp; *HSPA1B 3*′*UTR*, 379 bp; *PIN1 3*′*UTR*, 490 bp; *MAP2K2 3*′*UTR*, 277 bp) were amplified from HeLa genomic DNA and inserted between *NotI* and *XhoI* sites of the KDM6B-NU construct.

#### HP32Smut construct

In the KDM6B CDS corresponding to the IDR, 32/210 hydrophobic residues were mutated to serine. The DNA was synthesized by GenScript and cloned into the HA-tagged KDM6B construct using HiFi DNA Assembly (NEB). The sequence is shown in Table S8.

#### shRNA constructs

pLKO.1 puro was used to create shRNA constructs for stable knockdown cell lines. The sequence of the shRNA against KDM6A was previously reported^45^. shRNA sequences against ENO1 and DHPS were obtained from the RNAi Consortium at the Broad Institute. DNA oligonucleotides (Table S8) were annealed and inserted into pLKO.1 puro between *SgrAI* and *EcoRI*. As control, we used a scramble shRNA (Addgene #1864). To generate shRNA-resistant KDM6A expression constructs, eight synonymous mutations (CTATGAATCTCTAATCTT to tTAcGAgTCctTgATtcT) were introduced to the shRNA binding site of the original KDM6A plasmids using Q5 site-directed mutagenesis (NEB). shRNA-resistant DHPS and ENO1 expression constructs were generated with similar methods.

#### pET28a-MBP constructs

The pET28a vector containing a 6xHis-MBP tag was previously reported^12^ and used to clone all constructs for recombinant protein expression, except for the JMJD3 folded domain (FD) construct, which was cloned into the original pET28a backbone. For constructs used in in-vitro thiol-thiol crosslinking, a C-terminal Strep-Tag II (SAWSHPQFEK) was included in the reverse primer used to amplify the inserts. The JMJD3 FD (aa 1141-1643) were inserted between the *NheI* and *BamHI* sites. The amino acid boundaries of the JMJD3 regulatory IDR are aa 971-1178, whereas the regulatory IDR_RBRdel (RBRdel) aa boundaries are 1081-1178. To generate the 6xHis-MBP-reg_IDR-Strep-Tag-II construct, the wild-type regulatory IDR sequence was PCR-amplified and all the cysteines (C972, C990, C1138) were mutated to serines using the NEBuilder® HiFi DNA Assembly Master Mix (NEB, E2621L) with forward and reverse primers carrying the mutations and E1102 was mutated to cysteine for thiol-thiol crosslinking. As all native cysteines in the regulatory IDR were located outside of the HP region, they were deleted and a single cysteine was introduced in the HP region. According to AlphaFold3, E1102 is in close proximity to the folded domain in the predicted dimer. The 6xHis-MBP-reg_IDR_RBRdel-Strep-Tag-II construct was generated through PCR-amplification using the mutated regulatory IDR construct as template and inserted between the *NheI* and *BamHI* sites.

All constructs were verified by sequencing and all primer sequences and insert lengths are listed in Table S8.

#### DNA templates for RNA in vitro transcription

All DNA templates were PCR-amplified and purified by PCR cleanup. As the full-length *KDM6B* 3′UTR could not be transcribed, the sequence without the first 110 bp (1176 bp) was amplified from the pCMV-HA-KDM6B-U construct and used as *KDM6B* 3′UTR for in vitro experiments. The 3′UTR of HSPA1B (379 bp) was PCR-amplified from pCMV-HA-KDM6B-HSPA1B-UTR. The T7 promoter (TAATACGACTCACTATAGGG) was incorporated into the forward primers used for generating these templates.

### Plasmid transfection and viral transduction

Lipofectamine 3000 (Thermo Fisher, L3000015) was used to transfect pcDNA3-mGFP-MYC construct into U2OS cells. FuGENE HD (Promega, E2311) was used to transfect all pCMV-HA-derived plasmids and all pcDNA3-mGFP-KDM6B constructs into HeLa cells, except for HA-KDM6A, which was transfected into HeLa and MCF7 using Lipofectamine 3000. NU and U expression constructs for the same protein were transfected at the same molarity by default. For specific constructs requiring equalization of protein expression, the molarity was adjusted. Specifically, chimeric KDM6B constructs containing multivalent UTRs and U constructs of DHPS, SF3B4, SSRP1 and SSBP4 were transfected at a 1.3-fold higher molarity than their corresponding NU constructs. The FOXP2-U construct was transfected at a 1.5-fold higher molarity than its corresponding NU construct.

MCF7 cells stably expressing shKDM6B or HeLa cells stably expressing shDHPS or shENO1 were generated as previously described^11^. To generate virus, HEK293T cells were transfected using Lipofectamine 3000 with pCMV-dR8.2, pCMV-VSV-G, and the corresponding pLKO.1-shRNA plasmid. Viral particles were harvested 48 h after transfection and filtered through a 0.45 μm filter. MCF7 or HeLa cells grown in 6-well plates were transduced with 200 μl viral particles in the presence of 8 μg/ml polybrene. Puromycin (1 μg/ml) was added 48 h post-transduction. Cells were maintained in puromycin for at least four more days. Knockdown efficiency was validated by western blotting or or qRT-PCR.

### Western blotting

Cells were washed twice with PBS and lysed directly with 2x Laemmli sample buffer (Thermo Scientific, J61337.AC). The lysate was transferred to a 1.5 ml Eppendorf tube and boiled at 95°C for 10 min, followed by sonication for three times in 30s on/off cycles using a Diagenode Bioruptor. Samples were subjected to SDS-PAGE on NuPAGE 4 −12% Bis-Tris gradient gel (Invitrogen). For KDM6B, FOXP2, SSRP1 and KDM6A chemical crosslinking experiments, NuPAGE 3 - 8% Tris-Acetate protein gels were used. For thiol-thiol crosslinking *in vitro*, 6% or 8% Tris-Glycine gels were used. Membranes were scanned with an Odyssey DLx Imaging system (LI-COR). The primary antibodies used were: anti-GAPDH (1:2000, Sigma-Aldrich, G8795), anti-α-Tubulin (1:1000, Sigma-Aldrich, T9026), anti-Actin (1:1000, Sigma-Aldrich, A2066), anti-GFP (1:2000, Abcam, ab13970), anti-HA (1:2000, Covance, MMS-101P), anti-Hypusine (1:1000, Sigma-Aldrich, ABS1064-I), anti-EIF5A (1:1000, Cell Signaling Technology, 20765T), anti-ZAK (1:1000, Thermo Fisher, A301-993A), anti-TIAL1/TIAR (1:1000, BD Biosciences, 610352), anti-TIA1 (1:1000, Santa Cruz Biotechnology, sc-166247), anti-UTX (1:2000, Bethyl, A302-374A), anti-RPLP0 (1:1000, Antibodies Online, ABIN310003), and anti-KDM6B/JMJD3 (1:1000, abcam, ab169197). The secondary antibodies used were: IRDye 680RD donkey anti-chicken (LI-COR, 926-68075), IRDye 680RD goat anti-mouse (LI-COR, 926-68070), and IRDye 800CW donkey anti-rabbit (LI-COR, 926-32213).

### Chemical Crosslinking

Chemical crosslinking with DSG (Thermo Fisher) was performed as previous described^85^, with minor modifications. Briefly, HeLa cells were transfected with the indicated constructs. After 24 hours, cells grown in 24-well plates were washed twice with 0.5 ml PBS, and DSG in 200 μl PBS was directly added to each well. The reaction was incubated at room temperature (RT) with shaking for 30 min and quenched by adding 50 mM Tris, pH 7.5, for 15 min at RT. After discarding the buffers, cells were lysed with 2x Laemmli buffer. The samples were sonicated 3-5 times in 30s on/off cycles using a Diagenode Bioruptor and boiled at 95°C for 10 min. The crosslinked proteins were then analyzed by SDS-PAGE followed by western blotting.

For crosslinking using BS3 (Thermo Fisher) prior to mass spectrometry analysis, transfected cells from two 15-cm dishes were first lysed in a minimal buffer containing 20 mM HEPES, pH 7, 150 mM NaCl, 1% NP-40, and protease inhibitor cocktail. The lysates were triturated 20 times through a 26 Guage needle and cleared by centrifugation for 10 min at 14,000 rpm at 4°C. Crosslinking was performed on ice using 0.5 mM final BS3 and incubated for 2 h. The reaction was then quenched by Tris, and HA-tagged constructs were immunoprecipitated, eluted, and resolved on SDS-PAGE as described prior to being cut and submitted for mass spec analysis.

### Crosslinking mass spectrometry

#### Protein digestion and peptides desalting

Excised gel sections were washed with 100 µL of 1:1 (v/v) solution of 100 mM ammonium bicarbonate (ABC) and acetonitrile (ACN). After incubation for 15 minutes, the buffer was discarded, and 500 µL of ACN was added until the gel bands were destained. The gel pieces were then dried in a SpeedVac concentrator, reduced with 10 mM dithiothreitol (DTT), and incubated for 30 minutes at 56°C. Following reduction, 100 % ACN was added and incubated for 10 minutes at room temperature, then removed. Alkylation was performed by adding 55 mM iodoacetamide (IAA) and incubating the samples in the dark for 30 minutes at room temperature. After removal of the solution, the gel pieces were washed with 100 mM ABC and incubated for 10 minutes. The buffer was discarded, and 100% ACN was added twice for 5 minutes each. The gel pieces were then dried in a SpeedVac for 10 minutes and digested overnight at 37°C with shaking using 10 µL of mass spectrometry-grade trypsin at a concentration of 25 ng/µL in 50 mM ABC. Peptides were extracted using 5% formic acid (FA)/50% ACN and incubated for 15 minutes at 37°C with shaking. Peptides were dried in a SpeedVac and resuspended in 50 µL of 0.1% FA. Peptides desalting was performed using hand-packed stage tips containing Empore C18 extraction disks, which were sequentially conditioned with 100% methanol, 80% ACN/0.1% acetic acid, and 0.1% FA twice, prior to sample loading. After loading, the stage tips were washed with 0.1% FA. Peptides were eluted twice with 80% ACN/0.1% acetic acid, dried in a Speed-Vac centrifuge, reconstituted in 0.1% FA, water-bath sonicated, and transferred to liquid chromatography vials.

#### Mass spectrometry analysis

Peptides were separated on a 50 cm C18 column (Thermo Fisher ES903) using a gradient from 1.6% to 4% buffer B over 2 minutes, followed by an increase to 30% buffer B over 133 minutes, and then to 90% buffer B over 3 minutes, which was held at 90% buffer B for an additional 7 minutes (buffer A: 0.1% FA in HPLC-grade water; buffer B: ACN containing 0.1% FA). Separation was performed at a flow rate of 300 nL/min using a Vanquish Neo HPLC system (Thermo Fisher Scientific).

Mass spectrometry data were acquired on an Orbitrap Fusion Lumos mass spectrometer (Thermo Fisher Scientific) using a data-dependent acquisition (DDA) method with a 2 second cycle time. The method consisted of one MS1 scan at 60K resolution, an AGC target of 400,000, a maximum injection time of 50 ms, and a scan range of 380-1500 m/z. A charge state filter was applied to include charge states 4-8. MS2 scans were acquired using stepped collision energies of 21%, 26% and 31%, with a maximum injection time of 70 ms, an isolation window of 1.6 m/z, and a resolution of 30K.

#### Identification of crosslinked peptides

Peptides cross-linked by BS3 were identified by pLink3 software (version 3.0.16)^86^ using the following parameters: Cross-linker: BS3 (+138.068 Da); database: customized database with HA-JMJD3 sequence and reverse sequence added; enzyme: trypsin with up to 3 missed cleavages; precursor tolerance: 10 ppm; fragment tolerance: 20 ppm; fixed modification: carbamidomethyl [C] (+57.021 Da); variable modification: oxidation [M] (+15.995 Da) and deamidated [N] (+0.984 Da); filter tolerance: 10 ppm; separate false discovery rate (FDR) ≤5% at Peptide Pairs level.

#### Analysis of 3′UTR-dependent JMJD3 protein conformations

The KDM6B construct contains two proline insertions at aa 257, which were omitted in the analysis. Specifically, crosslinked residues were assigned based on the annotated JMJD3 sequence (UniProt ID: O15054-1) to one of the following JMJD3 regions: N-terminal IDR (aa 1-972), regulatory IDR (aa 973-1128), or folded domain (aa 1129-1682). Each crosslink was classified as IDR-IDR if either residue mapped within the N-terminal IDR or regulatory IDR region, otherwise the crosslink was classified as ‘with folded’. Crosslinks unique to individual samples (NU vs U) were visualized using xiVIEW^87^.

### Immunofluorescence staining

HeLa cells were seeded on 4-well Millicell EZ Slides and transfected. After 24 hours, the transfected cells were briefly washed twice with PBS, fixed with 4% paraformaldehyde for 10 min at RT, washed three times with PBS, and permeabilized in 0.1% Triton X-100 in PBS for 7 min. Cells were washed three times with PBST (PBS with 0.1% Tween-20) and incubated in blocking buffer (3% BSA in PBST) for 1 hour at RT. The cells were stained with the following primary antibodies diluted in blocking buffer at 4°C overnight: mouse anti-HA (1:1000, Covance, MMS-101P) and rabbit anti-H3K27me3 (1:2000, Cell Signaling Technology, 9733), or mouse anti-TIAR/TIAL1 (1:200, BD Biosciences, 610352). Cells were then washed 3x with PBST and incubated with secondary antibody in blocking buffer (goat anti-rabbit Alexa 647, 1:1000, Thermo Fisher A-21244 or donkey anti-rabbit Alexa 647, 1:1000, Thermo Fisher A-31575; donkey anti-mouse Alexa 488, 1:1000, Thermo Fisher A-21202) for 1 hour at RT. Cells were washed three times with PBST, mounted with mounting media with DAPI, and a coverslip was added. Samples were then imaged using a Nikon CSU-W1 SoRa spinning disk microscope with a 63x/1.40 CFI Plan Apo oil immersion objective. All images were taken at the same laser intensity and exposure settings. For endogenous TIAL1 in HeLa, images were acquired using a 4x magnification changer for SoRa and deconvolved with default settings using NIS-elements software.

### Lactate accumulation measurement

To measure the cellular activity of ENO1, shRNA-resistant HA-tagged ENO1 constructs were transfected into ENO1 knockdown or control HeLa cells. Twenty-four hours after transfection, the cells were quickly washed once with PBS, and the media replaced with DMEM, free of phenol red and FBS (Thermo Fisher, 21063029). The supernatant was harvested 30 minutes after media exchange. The lactate accumulation in the supernatant was measured using fluorometric lactate assay kit (Sigma-Aldrich, MAK570) following the manufacturer’s instructions.

### irCLIP to detect RNA-protein interactions in cells

To test the RNA-binding ability of JMJD3 produced from different constructs in cells, irCLIP was performed as previously described with slight modifications^54,55^. HeLa cells cultured in 10 cm dishes were transfected with the indicated HA-tagged KDM6B plasmids for 24 h. Cells were then washed with PBS once, crosslinked with 254 nm UV-C light (1500 J/m^2^), and lysed in 1 mL irCLIP lysis buffer (50 mM Tris pH 7.5, 150 mM NaCl, 1 mM EDTA, 1% Triton X-100, 0.1% SDS, 1X protease inhibitor). After sonication for 6 times in 30s on/off cycles, lysates were cleared by centrifugation at 5000 g for 10 min at 4°C. 1% of each sample was saved as input control. Each lysate sample was incubated with 30 μl HA magnetic beads (Thermo Fisher) for 2 h at 4°C with rotation. Beads were then sequentially washed with the following ice-cold buffers: twice with 1 ml irCLIP lysis buffer, twice with 1 ml high stringency buffer (20 mM Tris pH 7.5, 120 mM NaCl, 25 mM KCl, 5 mM EDTA, 1% Triton X-100, 1% sodium deoxycholate, 0.1% SDS), twice with 1 ml high salt buffer (20 mM Tris pH 7.5, 500 mM NaCl, 5 mM EDTA, 1% Triton X-100, 1% sodium deoxycholate), twice with 1 ml low salt buffer (20 mM Tris pH 7.5, 5 mM NaCl, 5 mM EDTA, 1% Triton X-100), and twice with 1 ml NT2 buffer (50 mM Tris pH 7.5, 150 mM NaCl, 1 mM MgCl_2_, 0.05% NP-40). 10% beads were then removed for immunoblot analysis, and the rest of the beads were resuspended in 30 μl NT2 buffer containing 0.5 μl RNase A (Thermo Fisher) and 15% PEG400 (Sigma-Aldrich) for on-bead RNase digestion at 30°C for 15 min with shaking (1200 rpm) in a Thermomixer. RNase digestion was quenched by the addition of 1 ml high stringency buffer, followed by two washes with 0.3 ml PNK wash buffer (50 mM Tris pH 7.0, 10 mM MgCl_2_), and then resuspended in 30 μl PNK dephosphorylation mix (1x PNK buffer (NEB), 0.5 μl SuperaseIn (Thermo Fisher), 1 μl T4 PNK (NEB), 4 μl PEG400). Dephosphorylation reactions were conducted at 37°C for 60 min with shaking (1200 rpm) in a Thermomixer. Dephosphorylation mix was removed, and beads were washed with 0.3 ml PNK wash buffer. For ligation of IRDye-800–conjugated oligo to RNA crosslinked to protein, beads were resuspended in 30 μl RNA ligation mix (1X RNA ligase I buffer (NEB), 1 μl RNA ligase I (NEB), 0.5 μl IRDye800-labeled oligonucleotide, 5 μl PEG400, and 0.5 μl SuperaseIn) and incubated for 16 hours at 16°C with shaking in Thermomixer (1200 rpm). Ligation mix was then removed, and beads were washed twice with 0.3 ml PNK wash buffer, prior to elution of RNA-protein complexes in 20 μl 1x LDS Buffer (Thermo Fisher) + 10% β-ME at 80°C for 10 min. 5 μl of eluates, as well as input controls, were then resolved by SDS-PAGE on 3-8% Tris-Acetate NuPAGE gels (Thermo Fisher) and transferred to nitrocellulose membranes. Fluorescent RNA-protein complexes in the eluates were visualized on a LiCOR Odyssey imager. The same membranes were then subjected to immunoblotting using anti-HA antibody.

### Digitonin RNA-FISH

RNA-FISH against GFP was performed before and after digitonin extraction of the cytosol, as previously described^11,13^. Briefly, HeLa cells were seeded on 4-well Millicell EZ Slide and transfected with GFP-fusion constructs. Fourteen to 16 hours after transfection, cells were incubated with 400 μl of digitonin solution (40 μg/ml digitonin, 150 mM NaCl, 20 mM HEPES pH 7.4, 0.2 mM EDTA, 2 mM DTT, 2 mM MgCl_2_) for 10 min at 4°C. The digitonin solution was aspirated, and coverslips were rinsed once with 400 μl PBS. For untreated samples, cells were briefly rinsed twice with PBS and subjected to the same RNA-FISH procedure described below. Specifically, cells were fixed with 4% paraformaldehyde for 10 min at RT, followed by two 5-minute washes with PBS and permeabilized using 1 ml of 70% ethanol in each well for 8 hours at 4°C. The slides were washed once with 1 ml wash buffer (2x SSC, 10% formamide) for 5 min at RT. A 200 μl hybridization mix containing Stellaris FISH Probes for eGFP with Quasar 670 Dye (Biosearchtech, VSMF-1015-5) was prepared and added to each well as previously reported. Slides were incubated at 37°C overnight, followed by two 30-minute washes with 1 ml pre-warmed wash buffer at 37°C. After a brief wash with PBST, slides were mounted with ProLong™ Gold Antifade Mountant with DAPI DNA stain (Thermo Fisher). Images were captured using a Nikon CSU-W1 SoRa spinning disk microscope with a 63x/1.40 CFI Plan Apo oil immersion objective. Excitations were performed sequentially using 405, 488, and 633 nm laser wavelengths. Eleven z-stacks were taken at 0.5 μm increments.

### Co-translational disome assay

#### Monosome and disome isolation

For isolation of monosomes and disomes, HeLa cells were plated on 15 cm dishes and transfected with HA-tagged constructs. Twenty-four hours after transfection, cells were washed with 10 ml 1x PBS supplemented with 100 μg/ml CHX (Sigma Aldrich) and 10 mM MgCl_2_. The PBS solution was aspirated, and 100 μl of ice-cold 5x lysis buffer (250 mM HEPES, pH 7.0, 50 mM MgCl_2_, 750 mM KCl, 5% NP-40, 50 mM DTT, 500 μg/mL CHX, 125 U/ml Turbo DNase (Invitrogen) and cOmplete EDTA-free protease inhibitor (Roche)) was added. The cells were scraped from the plate, and 1 unit/μl of RNase I (Ambion) was directly added to the lysate. The lysate was transferred to a 1.5 ml RNase-free Eppendorf tube and incubated on ice for 30 min.

The cell lysates were gently triturated ten times through a 26 Guage needle and cleared by centrifugation for 10 min at 14,000 rpm at 4°C. Cleared lysates were loaded onto 10-25% sucrose gradients in sucrose buffer (50 mM HEPES, pH 7.0, 5 mM MgCl_2_, 150 mM KCl, 100 μg/ml CHX, and EDTA-free protease inhibitor) and subjected to ultracentrifugation at 35,000 rpm in a TH-641 swinging bucket rotor (Thermo Fisher) for 3 hours at 4°C. Fourteen equal-volume fractions were collected using a Piston Gradient Fractionator (Biocomp) with continuous monitoring of RNA at UV260. Fractions 8-10 were pooled to obtain the monosome fraction and fractions 11-13 were pooled as the disome fraction.

#### Puromycylation and detection of puromycylated nascent proteins

To label and detect HA-tagged nascent chains in monosome and disome fractions, each fraction was incubated with 100 μM puromycin (Millipore) for 5 min at 37°C with shaking. 10% of puromycylated fractions were precipitated by TCA (TCI) and saved as input samples to adjust for total amount of ribosome loading. The remaining fractions were incubated with 20 μl of anti-HA magnetic beads (Thermo Fisher) for 2 hours at 4°C with constant tumbling. Beads were washed three times with 0.5 ml PBST and resuspended in 20 μl 2x Laemmli buffer. Immunoprecipitated puromycylated proteins were resolved on SDS-PAGE and probed using anti-puromycin antibody (1:1000, MilliporeSigma, MABE343) and anti-HA (1:2000, Covance, MMS-101P). To normalize for sample loading, total ribosome content in the input of each fraction was assessed by probing for RPLP0 using rabbit anti-RPLP0 (1:1000, Antibodies Online, ABIN310003). Band intensity of the HA-tagged nascent chain (puromycin signal co-localized with the HA signal) in each IP fraction was quantified by FIJI and normalized to the RPLP0 signal from the corresponding input. This yielded the normalized abundance of HA-tagged nascent chain in each fraction. The ratio of HA-tagged nascent chain in the disome over the monosome fraction was calculated and plotted.

#### Rule out ribosome collision by ZAK phosphorylation analysis

ZAK phosphorylation was analyzed using Phos-tag gels followed by western blotting as previously described^61^. Briefly, the cells were either irradiated at 1200 J/m^2^ (UV-C) in culture media, followed by recovery at 37°C for 30 min, or left untreated. The cells were rinsed with PBS once and lysed in lysis buffer (20 mM Tris-Cl, pH 8, 150 mM KCl, 15 mM MgCl_2_, 1% Triton X-100, 1 mM DTT) supplemented with phosphatase inhibitor cocktail (CST, 5870) and protease inhibitor cocktail (Roche). Lysates were incubated on ice for 30 minutes and cleared by centrifugation at 15,000 rpm for 10 min. A final 2x Laemmli loading buffer was added and boiled at 95°C for 5 min. The samples were resolved on 7% SDS-PAGE containing 10.7 μM Phos-tag (Wako, AAL-107) and 21.3 μM MnCl_2_. The gels were rinsed with transfer buffer containing 1 mM EDTA at room temperature for 10 min, followed by 10 min wash with the transfer buffer without EDTA, and transferred to PVDF membranes following manufacturer’s instructions. Membranes were blocked with 5% non-fat dry milk for 1 hour at room temperature, washed twice in PBST, and then probed with antibody.

#### iPSC to mesendoderm differentiation

Mesoderm and mesendoderm are the first few days during the GiWi cardiomyocyte differentiation protocol, which was used here^88^. Once the iPSCs reach 70% confluency, they were detached by Accutase and plated at 250000 cells per well in a 12-well under 10 µM Y27632 ROCK inhibitor (StemCell Technologies, 100-1044) with Stemflex medium. The media without ROCK inhibitor was changed every day for two more days until cells were 90% confluent. RPMI 1640 (Thermo Fisher Scientific, 11875093) with B27 minus insulin (Thermo Fisher Scientific, A1895601) and 10 µM CHIR99021 (StemCell Technologies, 100-1042) was added during the first 24 hours, followed by a 48-hour incubation of RPMI 1640/B27 minus insulin with 2 µM CHIR99021 to maintain Wnt signaling. Induction of markers was monitored by flow cytometry.

### Flow cytometry

Cells were collected at d0, d1, d2, d3 of the differentiation protocol. The cells were dissociated with 0.25% Trypsin (Thermo Fisher Scientific, 25200056) and neutralized by RPMI 1640 with 10% heat-inactivated fetal bovine serum. LIVE/DEAD™ Fixable Near IR staining (Thermo Fisher Scientific, L34994) was carried out to label the dead cells at 4°C for 30 minutes before 2% paraformaldehyde fixation at room temperature for 15 minutes. The cells were then permeabilized by 0.1% Triton X-100 prepared in FACS buffer (0.5% bovine serum albumin in PBS) at room temperature for 10 minutes and incubated with antibodies diluted in a FACS buffer at 4°C for 30 minutes. After washing out the unbound antibodies, the cells were resuspended in FACS buffer and passed through a 40-µm strainer to generate single cell suspension. Flow cytometry was performed on a CytoFLEX analyzer (Beckman Coulter) and the results were analyzed using FCS Express 7 software (De Novo Software). The antibodies used were: anti-Brachyury-APC (1:1000, R&D Systems, IC2085A), anti-EOMES-BV421 (1:1000, BD Bioscience, 567166), anti-KDM6B-FITC (1:1000, Novus, NBP1-06640F), and anti-GATA6-PE (1:5000, CST, 26452S).

### In vitro experiments

#### Protein purification

All expression constructs were transformed into BL21 *E. coli* (NEB) for protein expression. With the exception of the regulatory IDR, all proteins were purified using TALON resin under native conditions, followed by size exclusion chromatography. Specifically, a fresh colony was used to inoculate an overnight culture in 5 ml LB/kanamycin media and shaken vigorously at 250 rpm at 37°C. The overnight culture was diluted in 500 ml LB media containing 50 mg/l kanamycin and grown at 37°C until OD600 reached 0.6-0.8. Protein expression was induced by adding 0.5 mM IPTG, and the bacteria were cultured at 18°C overnight. Bacteria were harvested by centrifugation at 6,000 g for 15 min. The pellet was resuspended in 30 ml cold lysis buffer (50 mM sodium phosphate pH 7.4, 600 mM NaCl, 1 mM PMSF), supplemented with 10 mg lysozyme (Life Technologies), 1x Halt protease inhibitor cocktail (Thermo Fisher), and 1 mg DNase I (Roche). After incubation on ice for 30 min, bacteria were sonicated on ice for 3 min 30 seconds with on/off intervals of 1 and 3 s (Misonix S-4000). The lysate was cleared by centrifugation at 38,000 g at 4°C for 30 min.

5 ml TALON metal affinity resin (TaKaRa Bio) was equilibrated with five column volumes of native lysis buffer. The supernatant of cleared bacteria lysate was transferred to a new 50 ml Falcon tube and incubated with TALON resin with rotation at 4°C for 2 hours. The slurry was transferred into a gravity column and washed with 10 column volumes of native wash buffer (50 mM sodium phosphate pH 7.4, 600 mM NaCl, 10 mM imidazole). The protein was eluted with five column volumes of elution buffer (50 mM sodium phosphate pH 7.4, 600 mM NaCl, 200 mM imidazole).

The TALON-eluted protein was buffer-exchanged into protein storage buffer (25 mM Tris-Cl pH 7.4, 150 mM NaCl, 1 mM TCEP, 5% glycerol, and 0.01% Tween-20) using centrifugal filters with a 30 kDa cutoff (Pall) and concentrated to 1 ml. The concentrated protein was spun down at 20,000 g for 5 min at 4°C, and the supernatant was loaded onto a Superdex 200 Increase 10/300 column using the AKTA Purifier system (Cytiva). Peak fractions were collected, aliquoted, and snap-frozen in liquid nitrogen for storage at −80°C.

As the 6×His-MBP-StrepII-tagged regulatory IDR protein domain degrades on the size exclusion column, a two-step purification was performed. Specifically, the regulatory IDR was first purified using TALON resins under native conditions as described above. The eluted protein was concentrated to 10 ml using a 30 kDa cutoff centrifugal filter. The concentrated protein was passed through 2.5 ml of pre-equilibrated Strep-Tactin® 4Flow® high-capacity resin (IBA Lifesciences) twice. The protein-bound beads were then washed with 5 column volumes of PBS and eluted with 5 column volumes of PBS supplemented with 10 mM desthiobiotin (Sigma Aldrich). The eluted regulatory IDR was buffer-exchanged into the protein storage buffer by using a 10 kDa cutoff centrifugal filter, aliquoted, and snap-frozen in liquid nitrogen for storage at −80°C. Protein concentrations were determined by BCA assay (Thermo Fisher). The integrity of the purified protein was examined by SDS-PAGE and Coomassie blue staining.

#### RNA *in vitro* transcription

All RNAs were prepared by *in vitro* transcription using the T7 MEGAscript kit (Invitrogen). 250 ng of PCR-amplified DNA template was used for each 20 μl reaction. For Cy5 labeling, Cy5-UTP (Enzo life sciences, ENZ-42506) was added at a ratio of 1:300 labeled UTP:unlabeled UTP. The reaction mix was prepared following the manufacturer’s protocol and incubated at 30°C for 6 hours. The products were then treated with Turbo DNase provided in the kit and incubated at 37°C for 15 min. The integrity and sizes of the RNAs were verified using agarose gel electrophoresis. All RNAs were precipitated with LiCl overnight at - 20°C. The RNAs were pelleted by centrifugation at 20,000 g for 20 min at 4°C, washed with 1 ml of 70% ethanol, redissolved in nuclease-free water, aliquoted, and stored at −80 °C. RNA concentration was measured by NanoDrop.

#### Fluorescence polarization assay to measure RNA affinity

MBP-tagged protein domains were serially diluted using dilution buffer (25 mM Tris-Cl pH 7.4, 150 mM NaCl, 0.01% Tween-20, and 400 U/ml SUPERase·In™ RNase Inhibitor (ThermoFisher). Then, Cy5-labeled RNA (2.4 ng/µl) was added to the diluted proteins for a final reaction volume of 20 µL. All experiments were performed in triplicate. Reactions were incubated in a flat-bottom black 384 well-plate (Revvity) on a rocker at RT for 1 hour. Then, fluorescence polarization was measured using an Infinite M1000 Pro (Tecan) with the following parameters in i-control: excitation wavelength: 635 nm; emission wavelength: 665 nm; excitation/emission bandwidth: 5 nm; gain: 100 (manual); number of flashes: 30; settle time: 5 ms; Z-position: 24836 µm (manual); G-Factor: 1.1 (manual). Each sample was measured three-times, and the mean values were used for binding affinity calculations. Binding curves were fitted to the fluorescence polarization data using the nonlinear regression “One site – Total” model in Prism 10.

#### In vitro thiol-thiol crosslinking

To perform thiol-thiol crosslinking, JMJD3 regulatory IDR (5 µM) and FD (5 µM) were mixed with Cy5-labeled RNA (6 ng/µl) on ice in an assay buffer containing 25 mM Tris-Cl pH 7.4, 150 mM NaCl, and 0.01% Tween-20. BMOE (Thermo Fisher) was added to a final concentration of 5 µM in a total reaction volume of 20 µl. After 1 hour of incubation at 25°C, the reactions were quenched with 10 µl of 6x Laemmli SDS sample buffer. The samples were boiled at 95°C for 7 min and loaded onto 6% and 8% Tris-Glycine gels to resolve the dimer or monomer bands, respectively. Western blot was then performed, and proteins were probed with Streptavidin-680 (LI-COR, 1:3000 dilution).

### RNA-seq

To identify 3′UTR-dependent MYC targets, U2OS cells were transfected with GFP-MYC constructs and FACS-sorted 24 hours post-transfection to isolate GFP-positive cells. RNA was extracted from GFP-positive cells using TRIzol according to the manufacturer’s protocol and submitted to Genewiz for standard, 150 bp paired-end RNA sequencing (53-67 million reads were obtained per sample).

To identify 3′UTR-dependent KDM6A targets, endogenous *KDM6A* was knocked down with shRNAs and HA-tagged shRNA-resistant KDM6A constructs were transfected into the *KDM6A* knockdown cells. At 24 hours post-transfection, expression levels of total KDM6A and HA-tagged constructs were validated by western blotting using rabbit anti-UTX (1:2000, Bethyl, A302-374A) and mouse anti-HA (1:2000, Covance, MMS-101P) antibodies, respectively. RNA was then extracted using TRIzol according to the manufacturer’s protocol and submitted to Genewiz for standard, 150 bp paired-end RNA sequencing (19-30 million reads per sample were obtained).

### ChIP-seq

To analyze genome-wide 3′UTR-dependent JMJD3/KDM6B protein localization and activity, HeLa cells were co-transfected with HA-tagged KDM6B constructs and pcDNA3-mGFP at a 10:1 ratio. Twenty-four hours after transfection, cells were trypsinized, washed twice with ice-cold PBS, and stained with LIVE/DEAD fixable far-red stain (Thermo Fisher) following the manufacturer’s protocol. The cells were washed twice with ice-cold PBS and fixed with 1% paraformaldehyde for 10 min at room temperature on a nutator. Paraformaldehyde was quenched with 0.125 M glycine and mixed for 5 min at RT. The cells were centrifuged at 500 g for 5 min at 4°C, washed twice with ice-cold PBS supplemented with 1% FBS and EDTA-free protease inhibitor, and sorted by FACS to collect GFP-positive cells.

ChIP was performed by the MSKCC Epigenetics Research Innovation Lab. Cells were resuspended in SDS buffer (100 mM NaCl, 50 mM Tris-HCl pH 8.0, 5 mM EDTA, 0.5% SDS, 1x protease inhibitor cocktail from Roche). The resulting nuclei were spun down, resuspended in the immunoprecipitation buffer at 1 ml per 0.5 million cells (SDS buffer and Triton Dilution buffer (100 mM NaCl, 100 mM Tris HCl pH 8.0, 5 mM EDTA, 5% Triton X-100) mixed in 2:1 ratio with the addition of 1x protease inhibitor cocktail (Millipore Sigma, #11836170001) and processed on a Covaris E220 Focused-ultrasonicator to achieve an average fragment length of 200-300 bps with the following parameters: PIP=140, Duty Factor=5, CBP/Burst per sec=200, Time = 20 min. Chromatin concentrations were estimated using the Pierce™ BCA Protein Assay Kit (ThermoFisher Scientific #23227) according to the manufacturer’s instructions. The immunoprecipitation reactions were set up in 500 μl of the immunoprecipitation buffer in Protein LoBind tubes (Eppendorf, #22431081) and pre-cleared with 50 μl of Protein G Dynabeads (ThermoFisher Scientific #10004D) for two hours at 4°C. After pre-clearing, the samples were transferred into new Protein LoBind tubes and incubated overnight at 4°C with 5 μg of HA antibody (CST, #3724) and H3K27me3 antibody (CST, #9733). For normalization purposes, 2 μl of KMet SNAP-ChIP Panel (Epicypher, #191100) were added to each reaction. The next day, 50 μl of BSA-blocked Protein G Dynabeads were added to the reactions and incubated for 2 hours at 4°C. The beads were then washed twice with low-salt washing buffer (150 mM NaCl, 1% Triton X-100, 0.1% SDS, 2 mM EDTA, 20 mM Tris HCl pH 8.0), twice with high-salt washing buffer (500 mM NaCl, 1% Triton X-100, 0.1% SDS, 2 mM EDTA, 20 mM Tris HCl pH 8.0), twice with LiCL wash buffer (250 mM LiCl, 10 mM Tris HCl pH 8.0, 1 mM EDTA, 1% Na-Deoxycholate, 1% IGEPAL CA-630) and once with TE buffer (10 mM Tris HCl pH 8.0, 1 mM EDTA). Samples were then reverse-crosslinked overnight in the elution buffer (1% SDS, 0.1 M NaHCO3) and purified using the ChIP DNA Clean & Concentrator kit (Zymo Research, #D5205) following the manufacturer’s instructions. After quantification of the recovered DNA fragments, libraries were prepared using the ThruPLEX®DNA-Seq kit (R400676, Takara) following the manufacturer’s instructions, purified with SPRIselect magnetic beads (B23318, Beckman Coulter), and quantified using a Qubit Flex fluorometer (ThermoFisher Scientific) and profiled with a TapeStation (Agilent). The libraries were sent to MSKCC Integrated Genomics Operation core facility for sequencing on an Illumina NovaSeq 6000 (aiming for 30-40 million 100 bp paired-end reads per library).

### qRT-PCR

RNA was isolated from cells using TRI Reagent (Invitrogen) following the manufacturer’s protocol. RNA was reverse transcribed using qScript™ cDNA SuperMix (Quanta Bioscience). cDNA was diluted 5x and used for real-time PCR with gene-specific primers in the presence of PowerUp SYBR Green Master Mix (Life Technologies) by QuantStudio 6 Real-Time PCR system (Applied Biosystems). RPLP0 or GAPDH expression was used as housekeeping genes for normalization control. Primer sequences are listed in Table S8.

### Data analysis

#### mRNA features

The human MANE gene annotations (GRCH38/hg38) were used for all analyses^89^.

##### PhyloP sequence conservation in 3′UTRs

The genome-wide phylogenetic *P* value (phyloP conservation score) was obtained from the PHAST package for multiple alignments of 99 vertebrate genomes with the human genome in a form of a bigwig file. The genome-wide phyloP score was mapped to the 3′UTR boundaries of the human GTF (genome-build: GRCh38.p14, genome-build-accession: GCA_000001405.29), and the phyloP conservation values for 3′UTRs were obtained. To determine the number of conserved nucleotides in human 3′UTRs, we used a minimum phyloP score ≥ 2 and designated them as conserved, if at least four consecutive nucleotides had phyloP scores above this cutoff. If a given 3′UTR contains more than 150 conserved nucleotides, is it considered highly conserved. Non-conserved 3′UTRs contain zero conserved nucleotides, given our criteria. The sum of conserved nucleotides for each 3′UTR is listed in Table S1.

##### Enriched motifs in HC and NC 3′UTRs

To identify enriched 6-mer motifs in HC vs NC 3′UTRs, we used HOMER^90^. To identify motifs enriched in HC 3′UTRs, we used all HC 3′UTRs and compared them against the rest of the 3′UTRs. For motifs enriched in NC 3′UTRs, we used all NC 3′UTRs and compared them against the rest of the 3′UTRs. For all significant motifs, we calculated the motif frequencies within the groups of HC 3′UTRs and NC 3′UTRs by normalizing them by the total number of nucleotides in each group. Shown is the fold-change in motif frequency between HC and NC 3′UTRs.

##### mRNA features

3′UTR length, CDS length, and AU content were obtained from MANE and shown in Table S1.

##### Predicted RNA binding site density

As RNA valency describes accessible RNA-RNA binding sites in an RNA, we calculated the number of all possible 5-mer binding sites in a given RNA. We used all consecutive 5-mers and ask how often the reverse complement sequence is encountered in the RNA. We used Watson-Crick base pairing and allowed G-U wobble binding. We calculated the sum of all such ‘binding sites’ in a given 3′UTR. To normalize for RNA length, we divided this number by (3′UTR length)^2^ to obtain ‘binding site density’ or BSD. As this value does not account for accessibility of the putative binding sites, it describes RNA valency less well than A5/U5.

##### Mean MFE values of predicted RNA secondary structures obtained by RNAfold

We calculated the average minimum free energy (MFE) values for each 3′UTR obtained from RNAfold^91^. As MFE values depend on RNA length, we segmented each 3′UTR into 200-nucleotide windows with a 100-nucleotide overlap between consecutive windows. We used RNAfold to obtain the MFE values for each 200-nucleotide fragment and report the average MFE values obtained from all windows of a given 3′UTR in Table S1. Terminal 3′UTR windows shorter than 200 nucleotides were not included in the analysis.

##### A/U homopolymers (A5/U5)

Homopolymer stretches with at least five consecutive As or five consecutive Us, also called A5/U5, were counted in each 3′UTR and the sum is reported in Table S1. Ten consecutive As or Us were counted as two.

##### Determine the best surrogate marker for RNA valency

As RNA valency describes accessible RNA-RNA binding sites in an RNA^12^, Spearmen correlation coefficients between several parameters, including 3′UTR length, AU content, BSD, A5/U5, mean MFE, NED, condensate vs cytosol mRNA classification, and 3′UTR PhyloP conservation score were calculated (Table S5). This analysis revealed that A5/U5 correlated best with both 3′UTR length and weak RNA secondary structure (high mean MFE values). Therefore, A5/U5 was chosen as the surrogate marker for multivalent RNAs that strongly differ in length.

##### mRNAs enriched in cytosol or cytoplasmic network-like condensates

The list of cytosol-enriched mRNAs were obtained from Horste et al., Mol Cell (2023)^13^ and Chen et al., Cell (2024)^14^ and are listed in Table S1. CLIP peaks of condensate-enriched RNA-binding proteins were previously published^13^, as well as 3′UTR normalized ensemble diversity (NED) values^12^ and are listed in Table S1.

##### Codon usage

Translation rates can differ across codons. Based on two datasets^80,81^, we classified codons into optimal and non-optimal classes, and analyzed the ratio of non-optimal to optimal in the wildtype *KDM6B* CDS as well as in our mutants. We did not observe a difference in usage of non-optimal codons (Table S7).

### Protein features

#### Identification of IDRs

The coding sequences obtained from MANE gene annotations were translated into amino acids. A wrapper script was used to run IUPred2A^37^. All amino acids with IUPred2A score ≥ 0.33 were defined as IDRs and their number of amino acids are listed in Table S1.

To determine the longest IDR of each protein, we used a minimum of 30 amino acids and required ≥ 90% of amino acids to have IUPred2A values of ≥ 0.33. The longest continuous region that meets these criteria is annotated as longest IDR of a protein (Table S1). The longest folded domain of a given protein is calculated in a similar fashion, using a minimum of 30 amino acids and requiring ≥ 90% of amino acids to have IUPred2A values of < 0.33.

pLDDT (predicted Local Distance Difference Test) scores were obtained from AlphaFold and amino acids with pLDDT scores < 0.8 were considered unstructured. These values were used in Fig. 1G and S1B.

#### Hydrophobic amino acid clusters in IDRs (HP clusters)

Each longest IDR is scanned with a sliding window of four amino acids. If a window contains at least three hydrophobic amino acids (VILMCWYF), it is counted as hydrophobic sticker. An HP cluster is defined as at least two hydrophobic stickers with less than 20 amino acids of distance between them.

#### PhyloP sequence conservation in HP clusters and remaining IDRs

This analysis was performed for all MANE transcripts, whose CDS corresponds to uniprot amino acid sequence isoform 1.

The corresponding MANE coordinates of amino acid positions of HP clusters were determined and the average PhyloP scores within these regions as well as in the remaining IDR amino acids, present in the longest IDR were calculated.

#### Basic patches in IDRs

Basic patches are assigned to regions with window size of 10 with a net charge per residue ≥ 0.3. This means that within a stretch of 10 amino acids, we require four positively charged amino acids (K or R), if there is one negatively charged amino acid (E or D) present.

#### KDM6B net charge per residue (NCPR), hydrophobicity and PhyloP values

The net charge per residue for KDM6B was calculated with localCIDER^92^ using a step size of 50. Hydrophobicity values per residue were obtained from Protscale^93^. The PhyloP nucleotide scores for the KDM6B CDS were plotted with a sliding window of 50.

### Gene ontology analysis

Gene ontology analysis was performed using DAVID^39^. Bonferroni-adjusted *P* values are reported.

### Protein structure prediction

JMJD3 dimer structures were generated using the AlphaFold3 online server with default parameters. For each full-length protein, five structures were generated, and the highest-ranked structure was selected and rendered using UCSF ChimeraX (v1.9)^94^.

### RNA-seq analysis

Raw sequencing reads were quantified using Salmon (v1.10.1)^95^. The GENCODE release v44 and v47 basic annotations were used as the references for MYC RNA-seq and UTX RNA-seq, respectively. Transcript-level data was imported using tximport (v1.22.0)^96^, and differential gene expression analysis was performed using DESeq2 (v1.38.3 or Galaxy Version 2.11.40.8+galaxy0)^97^. Only protein-coding genes defined by the GENCODE annotation were included in the following analysis.

To identify genes regulated by MYC-NU or MYC-U, each condition was compared against the GFP-only control. We focused on mRNAs expressed in the GFP sample with mean TPM (transcripts per million) > 3. Differentially expressed genes have a |log2FC| > 0.3 and an adjusted *P* value < 0.1 in at least one of the tested conditions. Hierarchical clustering was performed on differentially expressed genes identified in either condition. The clustering was based on the z-score of the mean TPM value for each mRNA obtained from replicates and utilized the “ward.D” agglomeration method in R. 3′UTR-dependent groups were then defined using the obtained clusters and are presented in Table S2. The heatmap was visualized using the pheatmap (v1.0.13) package in R.

For the UTX RNA-seq analysis, UTX-regulated genes were first identified by comparing samples expressing shRNA targeting *KDM6A* to those expressing a scrambled control shRNA. Transcripts with a *P* value < 0.05 according to DESeq2 were considered significantly regulated. Based on the direction of change, these were classified as either “UTX-activated” (downregulated upon *KDM6A* knockdown) or “UTX-repressed” (upregulated upon *KDM6A* knockdown).

To identify genes rescued by expression of either the KDM6A-NU or KDM6A-U construct, we first filtered for robustly expressed genes, requiring a mean TPM > 7 in either the shCtrl or shKDM6A samples. We next selected genes that exhibited partial rescue, defined as those whose expression in KDM6A-NU or KDM6A-U transfected samples shifted back towards the levels observed in the shCtrl samples. mRNAs were divided into “UTX-activated” and “UTX-repressed” categories and clustered separately. Hierarchical clustering was performed on the z-scored mean TPM values using the “average” linkage method in R. 3′UTR-dependent groups were then defined using the clustering results and presented in Table S3.

### ChIP-seq analysis

Raw sequencing reads were trimmed and filtered for quality and library adapters using version 0.4.5 of TrimGalore (https://www.bioinformatics.babraham.ac.uk/projects/trim_galore), with a quality setting of 15 and running cutadapt (v1.15) and FastQC (v0.11.5). Reads were aligned to human assembly hg38 with version 2.3.4.1 of bowtie2 (http://bowtie-bio.sourceforge.net/bowtie2/index.shtml) and were deduplicated using MarkDuplicates in version 2.16.0 of Picard Tools^98^. To ascertain enriched regions, MACS2 (https://github.com/taoliu/MACS) was used with a p-value setting of 0.001 and run against a matched control for each condition. A peak atlas was created by combining the superset of all peaks using the ‘merge’ function in the BEDTools suite v2.29.2^99^. Read density profiles were created using deepTools ‘bamCoverage’ v3.3.0^100^, normalized to 10 million uniquely mapped reads and with read pileups extended to 200 bp. For histone modification data, ChIP signal was scaled to the inverse of the normalized counts to reads aligning to H3K27me3-modified synthetic nucleosome spike-in barcodes. H3K27me3 enrichment was classified as two-fold enrichment of spike in signal in comparison with control. Peak-gene associations were created by assigning all intragenic peaks to that gene, while intergenic peaks were assigned using linear genomic distance to transcription start site. Composite and tornado plots were created using deepTools v3.3.0 by running computeMatrix and plotHeatmap on normalized bigwigs with average signal sampled in 25 bp windows and flanking region defined by the surrounding 2 kb for HA signal and 10 kb for H3K27me3. Motif signatures were obtained using Homer v4.5 (http://homer.ucsd.edu).

### Immunofluorescence staining analysis

To analyze JMJD3/KDM6B protein abundance and protein activity after transfection of HA-tagged KDM6B constructs at the single-cell level, we loaded the obtained images using the nd2 (v0.10.0) package and processed them with the scikit-image package (v0.22.0)^101^. We first generated nuclear masks from DAPI channel grayscale images. Specifically, images were denoised using a Gaussian filter (sigma = 4, truncate = 4). Then, we created binary masks using each image’s mean intensity as a threshold. The masks were further refined through binary closing/opening, hole filling and removal of border-touching objects. Next, nuclei were segmented using the watershed algorithm, seeded by local maxima. Lastly, fluorescence signals from the H3K27me3 and HA channels were quantified using the refined masks.

The median fluorescence H3K27me3 signal obtained from untransfected cells (HA < 500 units) of each sample were used for normalization.

### Digitonin RNA-FISH analysis

To determine the fraction of mRNA transcripts retained in the cytoplasm after digitonin treatment, we calculated the ratio of mean fluorescence intensity in the cytoplasm of digitonin-treated cells over untreated cells. Specifically, we used FIJI^102^ to generate masks for the whole cell and nucleus using the maximum projection of fluorescence signals from RNA-FISH and DAPI channels, respectively. Next, maximum projection of RNA-FISH signals were quantified using the whole cell and nucleus masks. Mean cytoplasmic signals were calculated by subtracting total nuclear signals from total whole cell signals, divided by the cytoplasmic area. At least six fields of view were quantified for each sample from each replicate. For each experiment, mean cytoplasmic RNA-FISH signal in each digitonin-treated cell was divided by the median of the mean cytoplasmic RNA-FISH signal from all untreated cells. At least three independent experiments per construct were performed.

Amplicon seq analysis. Amplicon sequencing was performed by Genewiz. Paired-end reads were merged using Pear v0.9.6 with default settings^103^. Unique reads were counted with Seqkit^104^.

### Further statistical analysis

Statistical parameters are reported in the figures and figure legends, including the definitions and exact values of *N* and experimental measures (mean ± SD or SEM or boxplots depicting median, 25^th^ and 75^th^ percentile (boxes) and 5% and 95% confidence intervals (error bars)). All *P* values are listed in Table S9. Transcriptome-wide feature comparisons were performed using a two-sided Mann-Whitney test or a Wilcoxon test in SPSS. For experimental datat, *P* values were calculated using GraphPad Prism. Statistical analysis used unpaired, two-sided t-test for bulk analysis, and two-sided Mann-Whitney test for comparison of values obtained at single-cell levels.

## Notes

### Competing Interest Statement

The authors have declared no competing interest.

### Summary of Updates

Figures on JMJD3 crosslinking western blot removed; Data from partial deletion of endogenous KDM6B 3′UTR, JMJD3 crosslinking mass spectrometry, irCLIP, additional computational analysis, condensate-enriched RNA-binding protein, and additional protein candidates added, and corresponding text and method sections updated.

